# A multi-modal whole-slide image processing pipeline for quantitative mapping of tissue architecture, histopathology, and tissue microenvironment

**DOI:** 10.1101/2025.03.05.641761

**Authors:** Maomao Chen, Hongqiang Ma, Xuejiao Sun, Marc Schwartz, Randall E. Brand, Jianquan Xu, Dimitrios S. Gotsis, Phuong Nguyen, Beverley A. Moore, Lori Snyder, Rhonda M. Brand, Yang Liu

## Abstract

Multi-modal, multiscale imaging is crucial for quantitative high-content spatial profiling. We present an integrated image processing pipeline for comprehensive tissue analysis that combines quantitative phase microscopy for tissue architecture mapping, hyper-plex fluorescence imaging for immune microenvironment profiling, and whole-slide histopathology. This approach enables detailed morphological mapping of tissue architecture and cell morphology, while simultaneously linking them to the functional states of individual cells across the entire slide. By analyzing tissue biopsies from patients with ulcerative colitis, we demonstrate the potential of this pipeline for quantitative spatial analysis of molecular markers related to mucosal healing. Open-source and compatible with conventional microscopy systems, this pipeline provides a powerful tool for research and clinical applications through its comprehensive integration of quantitative, high-content, and histological imaging modalities.

## 1 Introduction

Spatial biology has recently emerged as a transformative field in biomedical research, offering valuable spatial context for the molecular characteristics of individual cells and providing deeper insights into their functional roles within the tissue microenvironment. While fluorescence imaging has been the primary method in spatial biology, visualizing fluorescently labeled targets, it focuses predominantly on selected molecular markers. To gain a more comprehensive understanding of tissue architecture, cell morphology, and physical properties—elements that are also diagnostically important [1, 2]—a multi-modal imaging approach is needed. This approach would integrate molecular profiling with unbiased structural and physical information, offering a more complete assessment of the tissue microenvironment and its cellular interactions.

Quantitative phase imaging (QPI) is a label-free technique that provides high-contrast images without external markers [2, 3], offering an unbiased view of tissue architecture and cell morphology [4], making it a valuable complement to fluorescence-based spatial molecular imaging, which maps molecular markers at sub-cellular resolution [5]. Additionally, histopathology remains the gold standard for clinical diagnosis and serves as a crucial reference for assessing pathological tissue states alongside these advanced imaging methods.

We present an integrated multi-modal image processing pipeline that aligns label-free QPI with fluorescence and H&E images, providing a comprehensive view of tissue architecture and function. To achieve high-resolution results, it incorporates multi-step resolution enhancement, aberration correction, and robust image stitching for whole-slide reconstruction. For hyperplex fluorescence imaging, it performs multi-cycle registration using the quantitative phase image as a reference, ensuring accurate alignment across imaging modalities and cycles. The entire process is fully automated and provided as open-source software, making it widely accessible for conventional microscopy systems.

To demonstrate the potential clinical utility of this multi-modal image processing pipeline, we conducted a pilot study on mucosal healing in patients with ulcerative colitis. The study quantitatively assessed tissue from patients during both active disease and remission, compared to healthy controls. The pipeline facilitated the analysis of nuclear architecture, cellular functions such as proliferation and immune activation, and the epigenetic states of epithelial cells and various immune cells. This approach demonstrates the feasibility of distinguishing between patients with active disease, those in remission, and healthy individuals, highlighting its potential for clinical applications in spatial biology and disease monitoring.

## 2 Results

### 2.1 Overall workflow of the image processing pipeline

**Figure 1** illustrates the general workflow of the automated image acquisition and processing pipeline. The workflow, described in detail in the Methods section, begins with the generation of quantitative phase images through phase retrieval from a series of bright-field images acquired at different axial positions, as shown in **Fig. 1(a)**. Next, a set of fluorescence images from different color channels are also acquired alongside bright-field images. We then computationally enhance the resolution and correct for aberrations (e.g., field of curvature aberration, spherical aberration) through region-based refocusing and deconvolution, as illustrated in **Fig. 1(b)**. To produce the whole-slide image, individual image tiles are captured at various locations across the slide, with approximately 10% overlap between adjacent images. These images are then stitched together, as shown in **Fig. 1(c)**, to produce whole-slide quantitative phase and fluorescence images. For hyperplex fluorescence imaging, we use multicycle fluorescence staining, imaging and bleaching/elution. To accurately align images from different imaging cycles, we use the unbiased tissue architecture from quantitative phase images as the reference for image registration. Lastly, **Fig. 1(d)** demonstrates the co-registration of multi-modal images, including quantitative phase image, hyperplex fluorescence image and whole-slide histopathology image from the same tissue slide. This integrated approach allows for comprehensive analysis of tissue morphology and molecular characteristics across different imaging modalities.

**Fig. 1.**
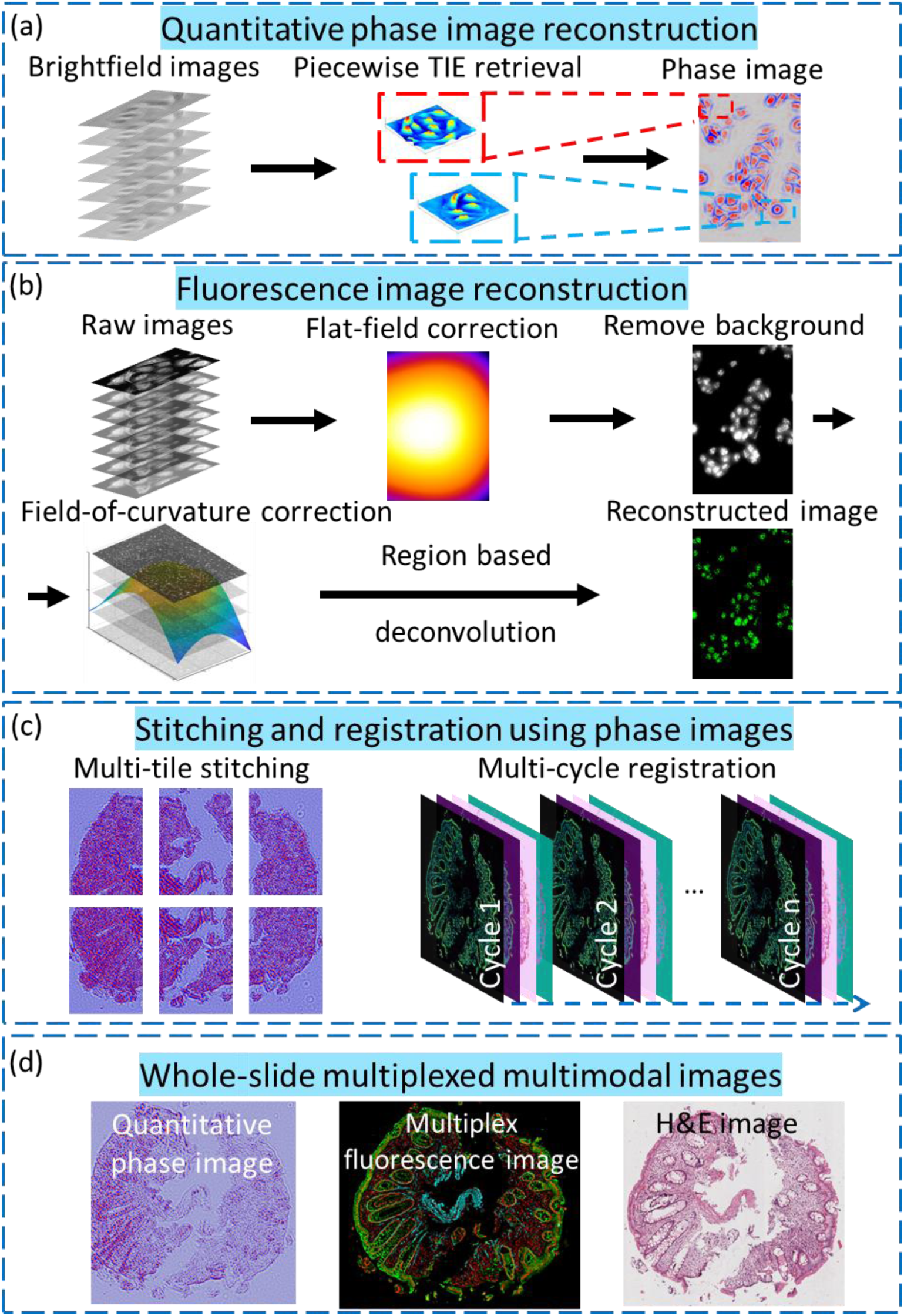
Overall workflow of the multi-modal image processing pipeline. **(a)** The reconstruction process of quantitative phase image based on the transport-of-intensity (TIE) equation. **(b)** The aberration correction and resolution enhancement of fluorescence image. **(c)** The whole-slide image is obtained by stitching multi-tile phase images acquired at different locations. The hyperplex fluorescence images are obtained by aligning multi-channel and multi-cycle images, and the transformation matrix is calculated using phase images. **(d)** The whole-slide multiplexed and multimodal images through multi-modal image registration.

### 2.2 Quantitative phase imaging using region-based transport-of-intensity equation

All microscopic imaging systems exhibit various aberrations, particularly field curvature aberration when imaging over an extended field of view (FOV). This results in defocused images at the edges of the FOV due to the curvature of the focal plane. To address this issue, we implement a region-based transport-of-intensity equation (TIE) phase retrieval method across the entire FOV.

As illustrated in **Fig. 2(a)**, we first divide the bright-field images into several smaller regions. Refocusing is applied to each region individually, and the quantitative phase image is reconstructed in each section by solving TIE. The reconstructed phase images provide detailed morphological information about the cells, as seen in **Fig. 2(a)**.

**Fig. 2.**
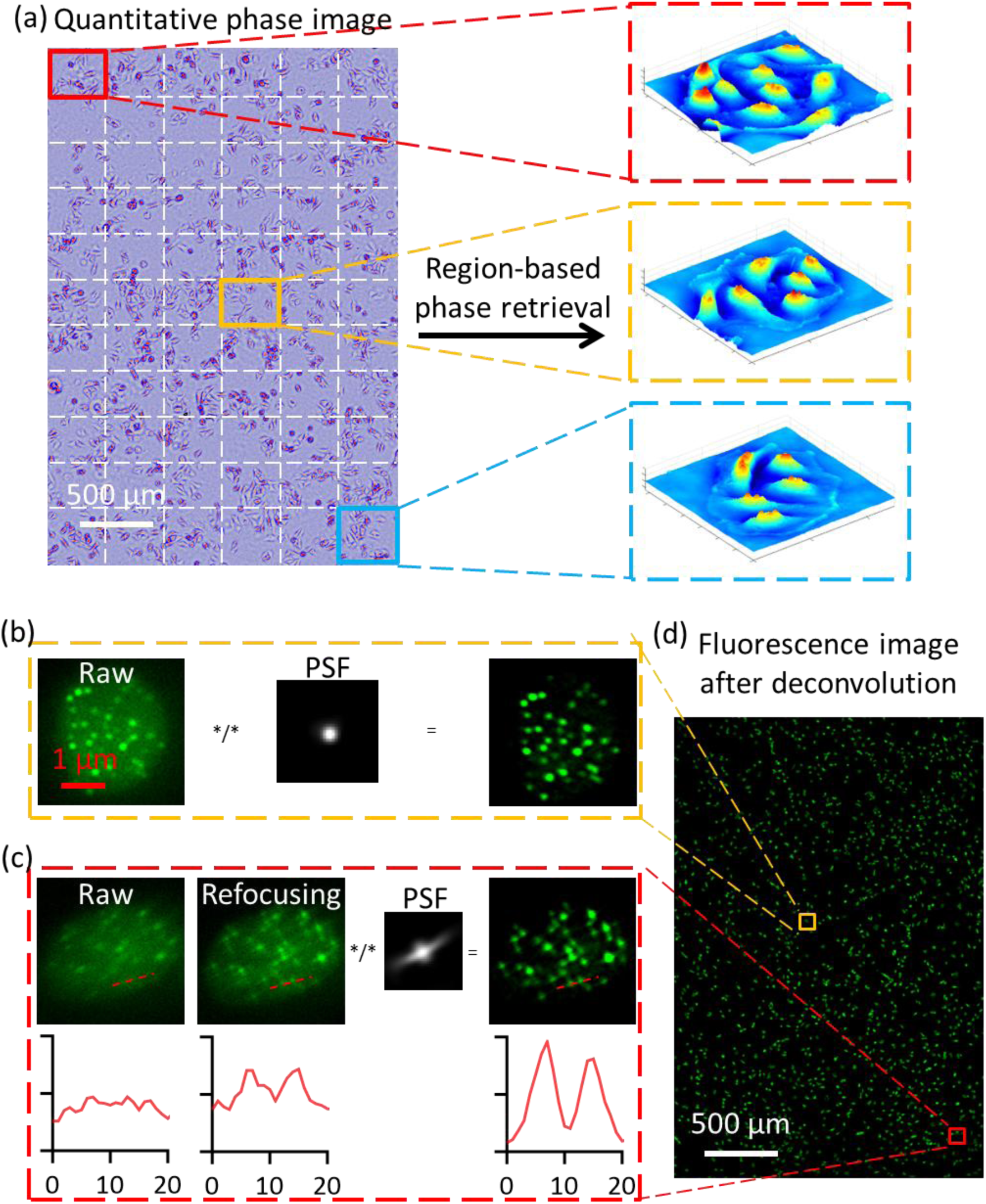
Region-based reconstruction of quantitative phase image and deconvolution of fluorescence image. **(a)** The bright-field image stack is divided into small regions, and the quantitative phase image of each region is obtained by region-based refocusing and TIE-based phase retrieval. **(b)**-**(c)** Refocusing and deconvolution of central (b) and corner (c) regions of the fluorescence images of the centromeres labeled against CENP-A. Sub-cellular structure of centromeres is clearly visible. **(d)** The improved fluorescence image after region-based refocusing and deconvolution.

**Supplementary Figure 1(a)** shows the final high-resolution phase image across the entire FOV, achieved by combining the phase images from all regions. **Supplementary Figure 1(b)** further demonstrates the improvement from region-based refocusing. When compared to traditional methods without refocusing (**Supplementary Fig. 1(b1)** and **(b3)**), our region-based TIE approach (**Supplementary Figs. 1(b2)** and **(b4)**) significantly enhances cellular details, especially at the corners of the image.

To evaluate the accuracy of the region-based TIE approach, we used 2 μm polystyrene microbeads embedded in glycerol, which have a theoretical phase value of approximately 2.96. **Supplementary Figure 2(a)** shows the quantitative phase image of the beads. **Supplementary Figure 2 (b)** compares the retrieved phase values of the beads at different regions, reconstructed using TIE-based phase retrieval with and without refocusing. While both methods under-estimate the retrieved phases, our region-based TIE method with refocusing shows a closer alignment with the theoretical estimation, with a more notable improvement at the corner regions of the image.

### 2.3 Flat-field correction and background removal of fluorescence images

Fluorescence images are acquired alongside bright-field images during the image acquisition process for hyperplex fluorescence imaging and high-content image analysis. To improve the accuracy and resolution of fluorescence quantification, we apply a series of computational enhancements, including flat-field correction, background removal, aberration correction, and deconvolution.

Uniformity in illumination and detection intensity is essential for intensity-based quantification of fluorescence signals. As shown in **Supplementary Fig. 3(a),** the non-uniformity causes region-based variation in the detected fluorescence intensity, which can bias the subsequent quantification. To correct for this system-induced uniformity across the entire image, we perform the flat-field correction by compensating the fluorescence images according to a calibration map as detailed in the Methods section.

Following flat-field correction, the fluorescence images show an improved uniformity across the FOV. To quantitatively assess this improvement, the intensity profiles along the diagonal black dashed line in **Supplementary Fig. 3(a)** are plotted in **Supplementary Fig. 3(b).** According to the curve fitted profile (red solid lines), the fluorescence intensity in the corner region improves from about 75% to 95% after flat-field correction, demonstrating the efficacy of the correction in achieving more consistent fluorescence signal across the image.

To further enhance the resolution of the fluorescence images, we apply a computationally efficient linear deconvolution approach. Although deconvolution is widely used to enhance the resolution of fluorescence images, it is susceptible to non-uniform background noise. As shown in **Supplementary Fig. 3(c)**, strong background noise can introduce artifacts, especially when the fluorescence signal from the target is weak. After background removal, as demonstrated in **Supplementary Fig. 3(d)**, the deconvolution significantly enhances contrast for sub-cellular cell structures, and the signal-to-noise ratio (SNR) is improved from 3.6 to 8.9.

### 2.4 Aberration correction and resolution enhancement of fluorescence images

To accurately estimate the point spread functions (PSFs) required for deconvolution, we experimentally determine the PSF across the entire FOV. As shown in **Supplementary Fig 4(a)**, the non-uniform patterns of PSF are evident due to the field curvature and spherical aberration, common in dry objective-based microscopy systems with an extended FOV. At the center of the image, the PSF appears Gaussian-like intensity distribution. However, as we move toward the edges of the field of view, the PSF becomes increasingly distorted.

To correct for region-dependent aberrations and achieve uniform resolution across the entire FOV, we apply a region-based deconvolution approach to fluorescence images, as shown in **Supplementary Fig 4(a)**. First, we refocus each region to accurately determine its focal plane. Next, we apply high-speed three-dimensional linear deconvolution [6, 7] using region-specific PSF. As shown in **Figs. 2(b-c)**, this method effectively corrects aberrations, particularly at the image edges, leading to significant improvements in both resolution and signal-to-noise ratio (SNR). The profile reveals that two centromeres, initially indistinguishable in the raw image, become clearly distinguishable after refocusing and deconvolution. The sub-cellular features of centromeres inside the nucleus are clearly visible in our imaging system. By applying this process to all regions, as demonstrated in **Fig. 2(d)**, we generate a fluorescence image with relatively uniform resolution across the entire FOV.

To quantitatively estimate the image resolution before and after processing, we applied the Fourier Ring Correlation (FRC)-based approach [8]. As shown in **Supplementary Fig. 4(c)**, the FRC resolution in the central region is improved from 1.3 μm to 0.7 μm, while the resolution in the peripheral regions is improved from 1.84 ± 0.13 μm to about 1 ± 0.17 μm. Overall, the fluorescence image exhibits approximately a 40% improvement in FRC resolution with our image processing pipeline.

### 2.5 Phase image-based stitching and registration

A mosaic scanning technique is often required to generate a whole-slide image, where different regions of the sample are scanned and then stitched together to form a complete image. While this method is well-established, it can present challenges when fluorescently labeled targets have weak signals or exhibit sparse structures, limiting the mutual information between adjacent image tiles and complicating accurate image registration.

To enhance the robustness of image registration and stitching, we leverage quantitative phase images, which provide unbiased and detailed architectural information. The shifts between adjacent image tiles are initially corrected using the phase correlation method [9, 10]. Once aligned, the large FOV image is constructed by fusing the individual tiles through a linear blending technique [10, 11].

**Figure 3(a)** shows a whole-slide image of a 6 mm x 5 mm tissue sample. After correcting for positional shifts between adjacent image tiles, the normalized cross-correlation coefficient (NCC) between neighboring images improves from 0.07 to 0.92, revealing clear glandular structures in the colon tissue without any visible discontinuities. The same transformation matrix is subsequently applied to all fluorescence channels, enabling the generation of whole-slide images for each fluorescence modality.

**Fig. 3.**
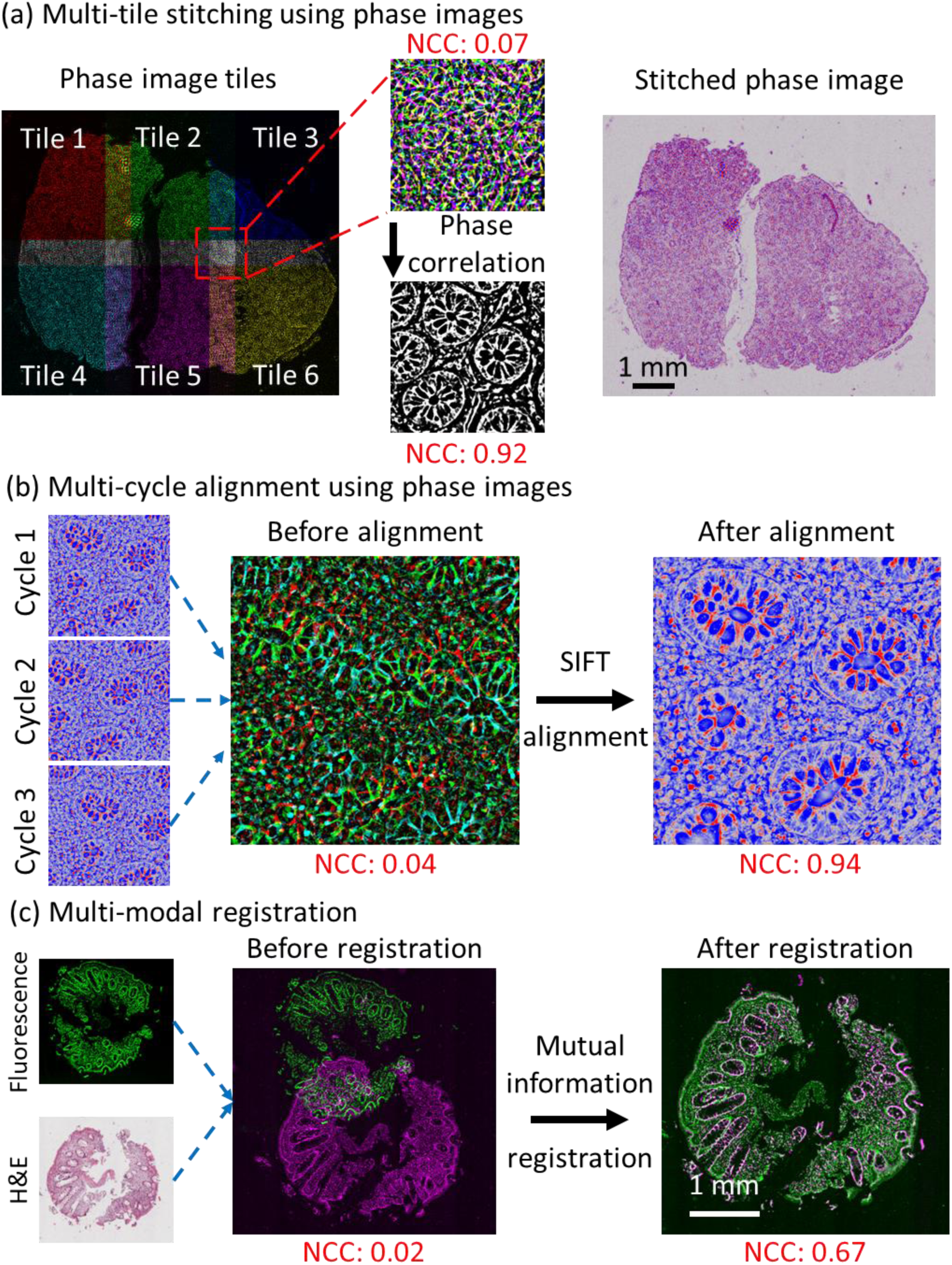
Image stitching and registration. **(a)** Multi-tile image stitching using the quantitative phase images as the reference. NCC: normalized cross-correlation coefficient. NCC: normalized cross-correlation coefficient. **(b)** Image alignment using quantitative phase images from multiple imaging cycles. **(c)** Multi-modal registration between the fluorescence and H&E-stained whole-slide histology images.

Building on the alignment process described earlier, cyclic immunofluorescence staining and imaging are frequently used to visualize multiple molecular markers (more than five) on the same tissue section for high-content profiling [12]. This technique typically requires the registration of fluorescence images from each imaging cycle. Traditional multi-cycle registration relies on a reference fluorescence marker, such as DAPI, in each cycle. With phase images for image alignment instead, our method eliminates the need for a reference marker, freeing up an imaging channel and improving overall efficiency.

As illustrated in **Fig. 3(b)**, the consistent tissue architecture across all channels and cycles allows us to eliminate the need for a dedicated fluorescence reference channel. This consistency streamlines the entire registration process and enhances the overall efficiency of multi-cycle imaging. We achieve alignment of the multi-cycle phase images using the Scale-Invariant Feature Transform (SIFT) method [13–15], effectively correcting misalignments between phase images from different cycles. The NCC improves significantly from 0.04 to 0.94. By applying the alignment information obtained from the phase images to the corresponding fluorescence images, we successfully align all multi-channel fluorescence data.

In addition to quantitative phase imaging and multiplexed fluorescence, Hematoxylin and Eosin (H&E)-based histopathology remains the clinical gold standard for diagnosis. The integration of histological and fluorescence images can be achieved using a mutual information-based registration method [16], as shown in **Fig. 3(c)**. This method aligns the images on a pixel-by-pixel basis [17, 18], ensuring precise correlation between the different imaging modalities. Such integration enables the visualization of multimodal information, providing a comprehensive understanding of the tissue microenvironment while preserving the context of traditional morphological assessments.

### 2.6 Multi-modal multi-scale images from an inflamed colon tissue

**Figure 4(a)** presents the co-registered whole-slide histology, quantitative phase, and hyperplex fluorescence images of inflamed tissue from a patient with ulcerative colitis. These multi-modal whole-slide images on the same tissue offer complementary insights into tissue architecture, morphology, and molecular characteristics at sub-cellular resolution. The quantitative phase image reveals overall tissue architecture, extracellular matrix, and cell morphology via the properties of refractive index. The hyperplex fluorescence images, visualizing multiple molecular targets, reveal the spatial distribution of various cell types along with their functional states in the context of the tissue architecture. Meanwhile, the H&E-stained histology image serves as the gold standard for conventional pathology assessment, offering a benchmark for comparison with the other modalities.

**Fig. 4.**
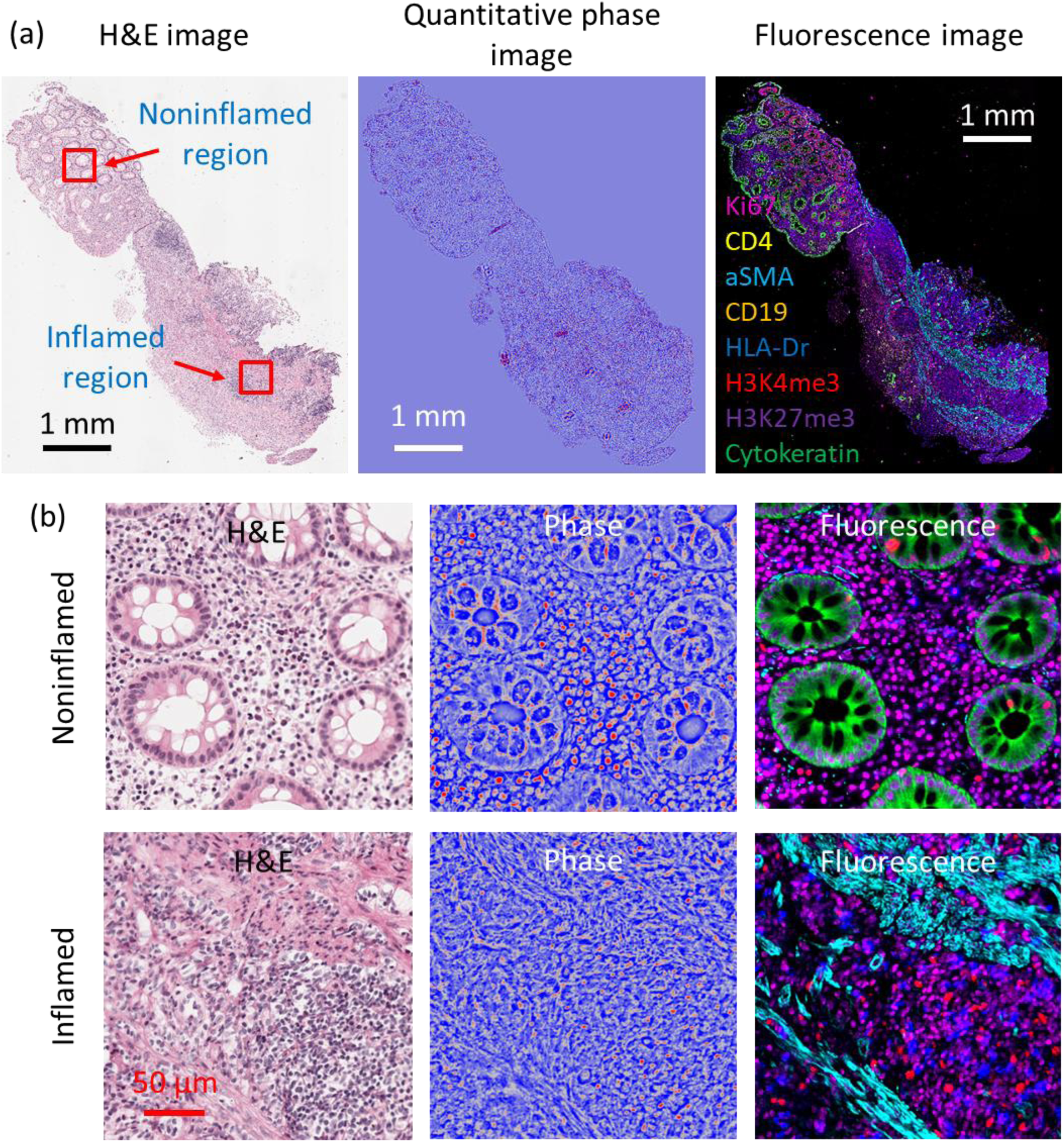
Multi-modal imaging of an inflamed tissue. **(a)** The H&E, quantitative phase and fluorescence images of the UC tissue. **(b)** The zoomed images of the non-inflamed region and the inflamed region of the tissue.

**Figure 4(b)** shows the structural differences between non-inflamed and inflamed areas of the same tissue. In the non-inflamed region, clear and intact glandular structures are observed, whereas the inflamed area is characterized by severe inflammation, dominated by immune cell infiltrates, disrupted stroma, and few remaining epithelial regions. The phase images further differentiate these areas, revealing that immune cells in the non-inflamed stroma have a higher optical density or refractive index compared to those in the inflamed region, indicating higher chromatin density in the inactive immune cells. Additionally, the label-free phase image clearly delineates the extracellular matrix in the inflamed area, providing further insight into the tissue’s structural changes during inflammation.

**Figure 5** presents an example of the whole-slide hyerplex fluorescence image from 16 protein targets on the same tissue. The images are obtained from four-cycle immunofluorescence staining, each comprising four distinct color channels. **Figure 5(a)** is a multiplexed image showing unique fluorescent channels for 8 of the 16 markers. **Figures 5(b-e)** provide examples of images produced from each staining cycle utilizing 4 markers per cycle. These highly multiplexed images enable the simultaneous visualization of 16 fluorescence markers, allowing for direct observation and analysis of various cell types and their functional states within the same tissue. The image reveals a densely populated area of immune cells within the inflamed region. In the area of severe inflammation, immune cells dominate, with minimal presence of epithelial cells. Activated immune cells, such as B cells (CD19+) and helper T cells (CD4+), show high expression of HLA-DR and TNFα, indicating an active immune response. In the images of cycle 3, the smooth muscle actin (αSMA) is prominently visible in the inflamed region, suggesting significant tissue remodeling. In contrast, the images from cycle 4 distinctly highlight the intact glandular structures and epithelial cells in the non-inflamed areas of the tissue, providing a clear contrast between inflamed and normal-appearing regions.

**Fig. 5.**
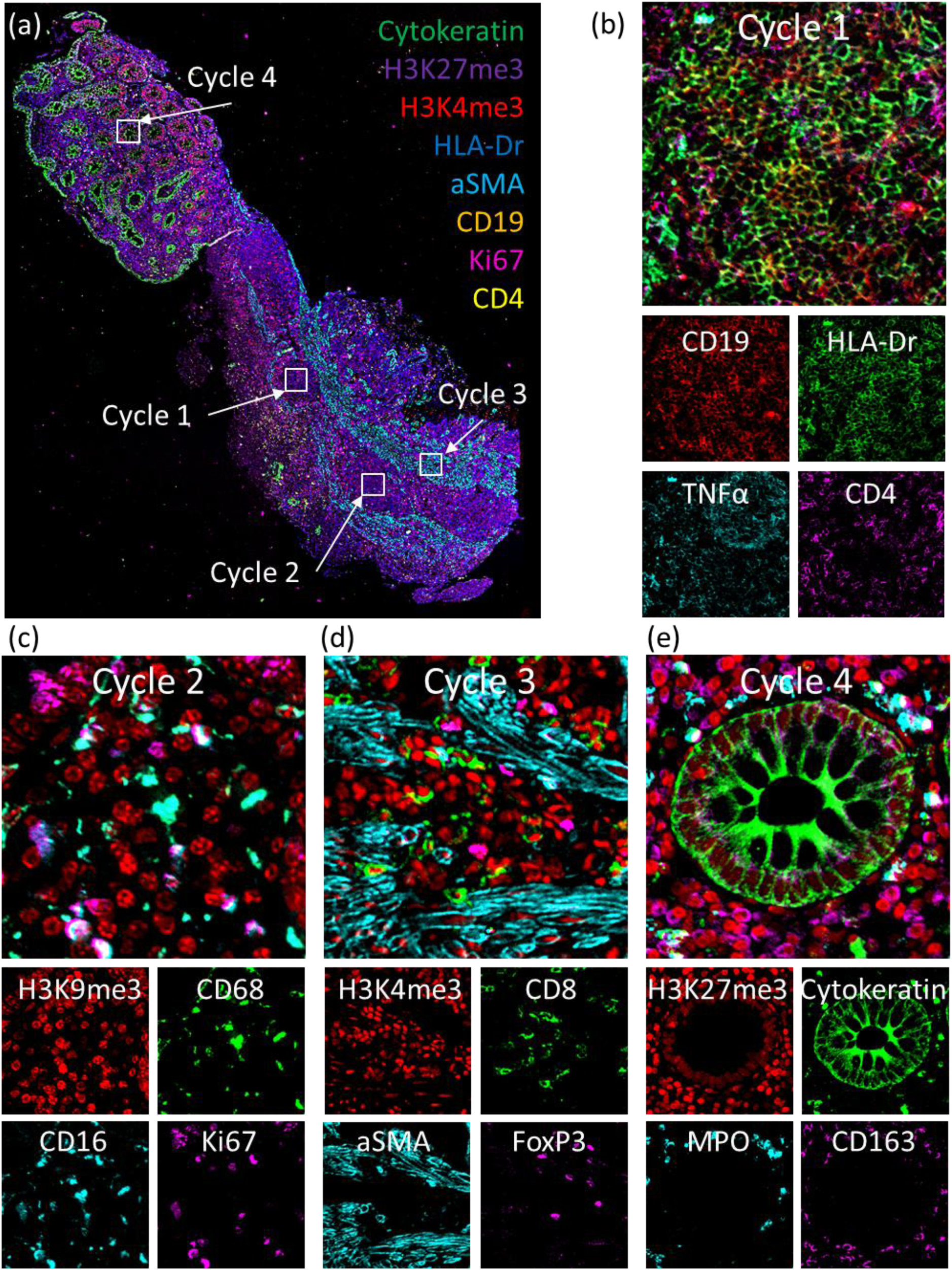
Hyperplex fluorescence images of the inflamed tissue. The multiplexed fluorescence images were obtained through 4 cycles of immunofluorescence staining and imaging with 16 protein markers.

### 2.7 Quantitative spatial analysis of cell types and functional states in mucosal healing

We quantitively evaluated mucosal healing in UC by analyzing distribution of epithelial and immune cell types, as well as their functional states, in tissue samples from three different groups of patients: normal tissue from healthy individuals, inflamed tissue from UC patients with active inflammation, and normal-appearing tissue from UC patients in remission. **Figure 6(a)** shows representative hyperplex fluorescence images from each group. In the control group (healthy individuals), we observed well-defined gland structures with relatively uniform shape and size. In contrast, the inflamed tissue from UC patients showed a notable disruption of gland structures, with a predominance of immune cells. Interestingly, the normal-appearing tissue from patients in remission exhibited partial recovery of gland structures, suggesting a healing process from UC. **Figure 6(b)** shows the corresponding spatial distribution of color-coded cell types in the form of Voronoi polygons. In healthy tissue, distinct and compartmentalized cell types are observed, while inflamed tissue from UC patients demonstrates a significant reduction in epithelial cells and a notable increase in immune cells, with highly inter-mixed cell populations. In remission, an intermediate status of mucosal healing is apparent, with partially restored, compartmentalized glands and a marked increase in neutrophils, reflecting the tissue’s ongoing recovery process.

**Fig. 6.**
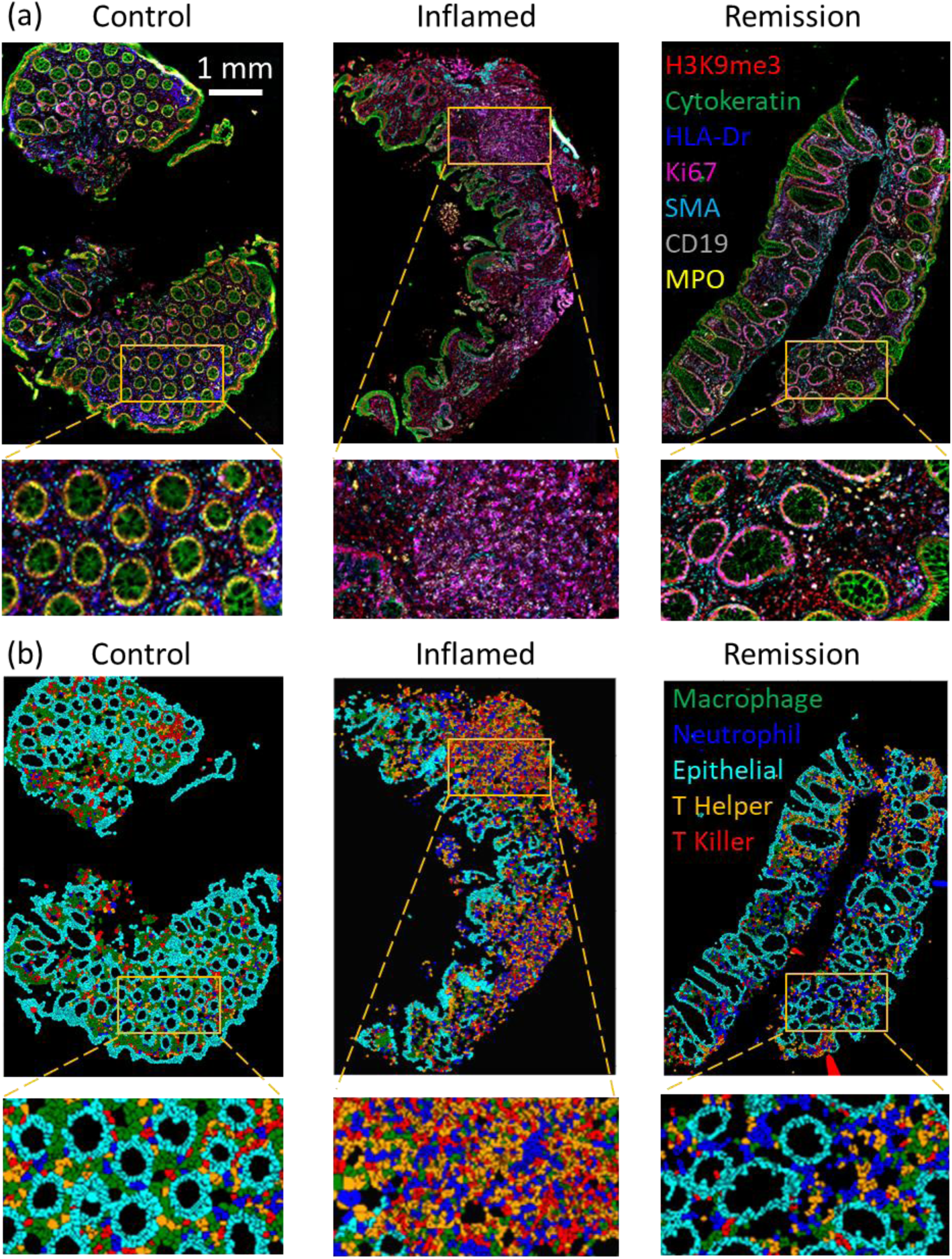
Fluorescence images and cell phenotyping. **(a)** The representative hyperplex fluorescence images of tissue samples from control, inflamed, and remission patients. **(b)** Spatial distribution of color-coded cell types in the form of Voronoi polygons.

Furthermore, **Figure 7** illustrates the distribution of major cell types across patients with active inflammation, remission and healthy controls. In the healthy controls, epithelial cells constitute approximately 50% (ranging from 44% to 54%), while main immune cell population—including helper T cells, macrophages, neutrophils, and cytotoxic T cells—make up around 40%. In contrast, inflamed tissues show a marked reduction in epithelial cells, averaging around 17% (ranging from 9% to 26%), accompanied by a significant increase in immune cells. Notably, the proportion of helper T cells rises from an average of about 11% in healthy controls to approximately 35% in inflamed tissues, while neutrophils increase from around 7% to about 17%. These elevated immune cell populations indicate a strong immune response in inflamed tissues. Tissues in remission display characteristics that fall between those of healthy and inflamed tissues. The percentage of epithelial cells in remission averages around 40%, significantly higher than in inflamed tissues, suggesting tissue recovery. However, the still-elevated presence of immune cells points to ongoing immune activity, even as the tissue transitions toward a more normal state compared to healthy controls.

**Fig. 7.**
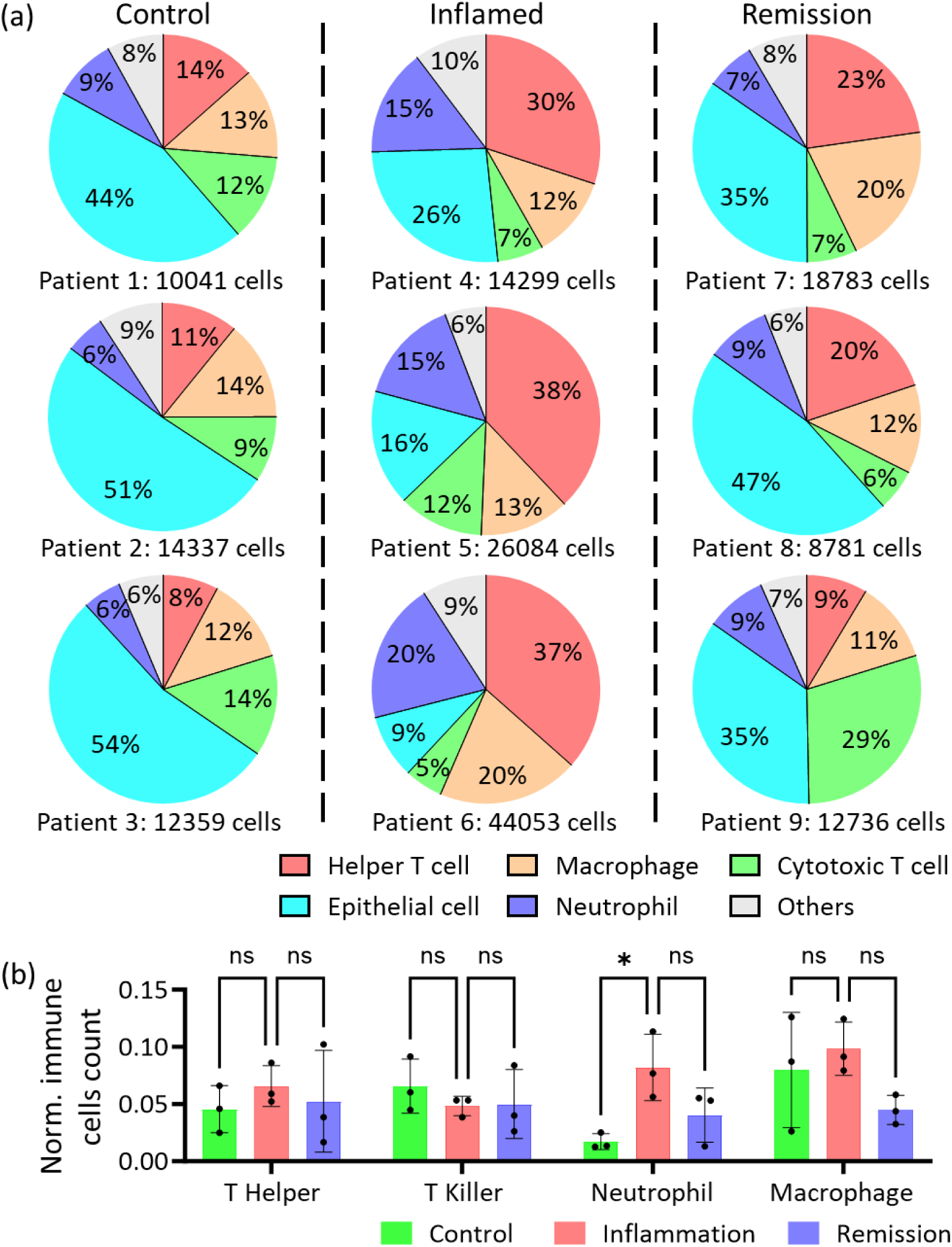
The percentage and the spatial adjacency of major cell types. **(a)** The percentage distribution of major cell types from colon tissue of healthy control patients and ulcerative colitis patients with inflamed and remission. Each pie chart shows the percentage of major immune and epithelial cell types from each patient. **(b)** The normalized adjacent immune cells around the epithelial cells (number of the adjacent immune cells divided by the number of epithelial cells). Data are presented as means ± standard deviation, and the statistical significance were determined using the two-way ANOVA test (*: *P*-value < 0.01; **: *P*-value < 0.001; ***: *P*-value < 0.0001; ns: no significance).

Additionally, **Figure 7(b)** presents a spatial adjacency analysis between epithelial and different types of immune cells. Although the small cohort limits the extent of our findings, the most significant spatial relationship observed is the high percentage of neutrophils located in close proximity to epithelial cells in inflamed tissues, as well as in some patients in remission. This proximity highlights the interaction between immune and epithelial cells during inflammation and ongoing mucosal healing.

**Figure 8** presents an analysis of the functional states of epithelial and immune cells. Here we focus on the commonly used immune cell activity and proliferation markers, including HLA-DR and Ki67. **Figure 8** reveals the percentage of active (HLA-DR positive) immune cells among different cell types in the three groups of patients. In patients with active inflammation, for all major immune subtypes, over 50% of immune cells are activated. Both neutrophils and macrophages showed the highest percentage of the active state. In contrast, less than 20% of immune cells (all major types) are in active states in healthy control tissues. Most importantly, in remission, our data demonstrates that the percentage of active immune cells falls between healthy controls and active inflammation, suggesting the importance of quantitative characterization in defining mucosal healing status. Similar to the inflamed tissue, macrophages, and neutrophils showed the highest percentage of active immune cells.

**Fig. 8.**
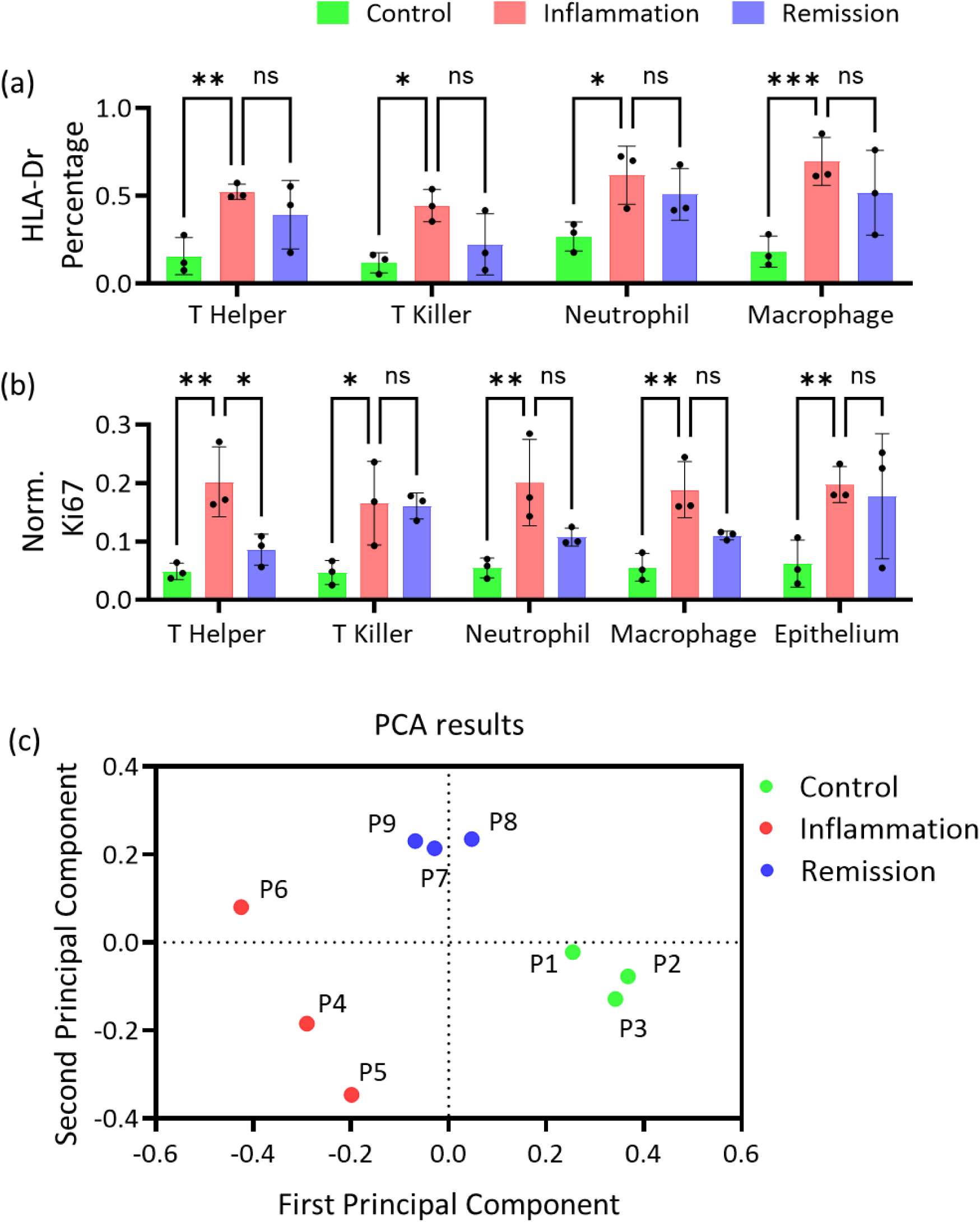
Functional analysis of the immune cells. **(a)** The percentage of the activated immune cells labeled with HLA-Dr marker. **(b)** The intensity of Ki67 of different immune cells. Data are presented as means ± standard deviation, and the statistical significance were determined using the two-way ANOVA test (*: *P*-value < 0.01; **: *P*-value < 0.001; ***: *P*-value < 0.0001; ns: no significance). **(c)** The separation of patient groups using principal component analysis. P: Patient.

Similarly, the mucosal healing status is also reflected in the cell proliferation marker Ki67. As shown in **Fig. 8(b)**, the percentage of Ki67-positive cells, including epithelial and the major types of immune cells, is significantly higher than those from normal-appearing tissue from healthy controls. This agrees with the well-established notion that inflamed tissue has greater cell proliferation. The percentage of Ki-67-positive cells for almost all major cell types is in an intermediate state between normal and inflamed cases. Interestingly, in patients in remission, we observed a greater level of heterogeneity among different cell types and patients. The cytotoxic T cells and epithelial cells also exhibit the highest percentage of Ki67-positive cells. These results suggest that in remission, despite being recovered from inflammation, there is still sustained immune response with a higher proportion of active immune cells in remission tissues and more proliferating cells.

Using various physical and functional parameters, patients were classified into three distinct groups using principal component analysis (PCA). As illustrated in **Fig. 8(c)**, the inflammation group displays a noticeably broader and more scattered distribution compared to the control and remission groups. This result highlights the greater biological variability within the inflammation group, consistent with clinical observations of UC heterogeneity reported by gastroenterologists.

### 2.8 Quantitative spatial analysis of cell types and functional states in mucosal healing

To accurately assess the molecular status of mucosal healing, the above two commonly used markers for assessing immune cell activation (HLA-DR) and cell proliferation (Ki67) are likely not sufficient for a larger patient cohort, due to the well-known heterogeneity of UC and the potential heterogeneous types of immune cell populations involved. Therefore, we examined a set of markers that reflect the global epigenomic states of the major cell types, focusing on chromatin compaction and repressive chromatin states, which may better capture immune cell activity while being less affected by patient heterogeneity.

First, the phase value reflects the refractive index of the cell nuclei, which is measured from label-free quantitative phase images. The phase values from **Fig. 9(a)** show that immune cells in inflamed tissue have a lower refractive index (RI) than those in control and remission tissue. This difference in refractive index largely reflects the changes in chromatin compaction, where the active immune cells exhibit an open chromatin state as we previously showed [19]. Therefore, this result suggests that inflamed cell nuclei, in epithelial and different types of immune cells, assume more open chromatin structure. This observation is further supported by examining the intensity of two histone markers, H3K27me3 and H3K9me3, as seen in **Figs. 9(b-c)**. These markers are typically associated with heterochromatin, which suppresses transcriptional activity. The observed lower intensity of H3K27me3 and H3K9me3 in the inflamed tissue suggests less condensed chromatin states, pointing to higher transcriptional activity in these cells.

**Fig. 9.**
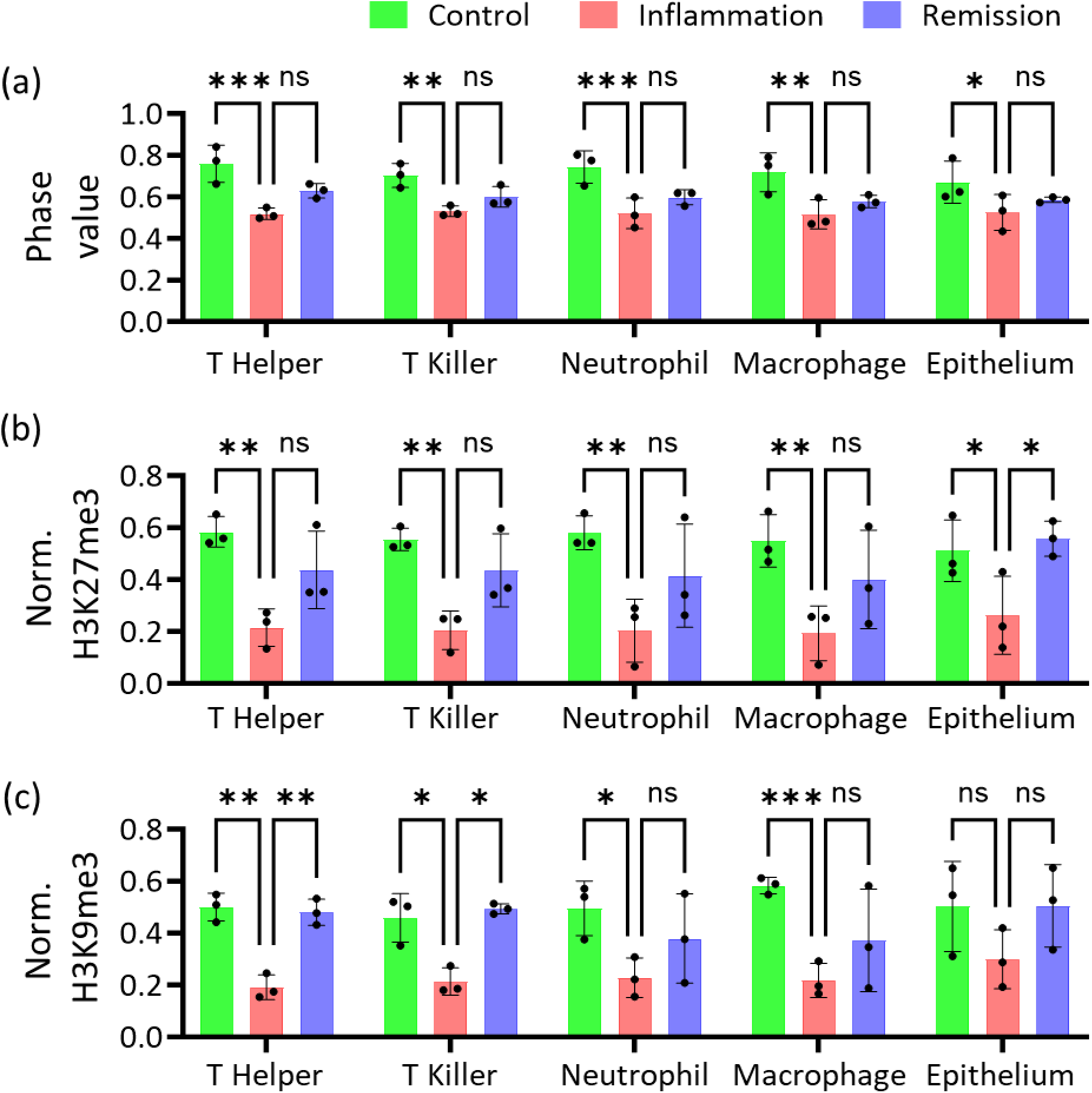
The mapping between the quantitative phase values and the histone structures of the immune cells. **(a)** The average phase values of the immune cells. **(b)** The normalized fluorescence intensity of H3K27me3 of the immune cells. **(c)** The normalized fluorescence intensity of H3K9me3 of the immune cells. Data are presented as means ± standard deviation, and the statistical significance was determined using the two-way ANOVA test (*: *P*-value < 0.01; **: *P*-value < 0.001; ***: *P*-value < 0.0001; ns: no significance).

## 3 Discussion

We have developed a multi-modal whole-slide image processing pipeline that integrates and interprets data from various imaging modalities, including label-free quantitative phase images for unbiased visualization of tissue architecture and hyperplex fluorescence images for high-content molecular characterization. This multi-modal integration provides a comprehensive view of both molecular and morphological features on the same sample, evaluated alongside the gold standard of histopathology. We provide the entire workflow for obtaining multi-modal high-resolution whole-slide quantitative phase and fluorescence images, encompassing phase retrieval from bright-field images, aberration correction, flat-field correction and resolution enhancement, mosaic image tile stitching and multi-cycle whole-slide image registration. By correlating fluorescence-based markers with phase-derived structural information and conventional histopathology, our platform offers deeper insights into tissue microenvironments and sets the stage for future AI-driven analyses across multiple imaging modalities. This image processing pipeline is offered as open-source software to the scientific community and can be widely used in laboratories with conventional light microscopy coupled with scanning stages, which is also commercially available as accessories.

Technically, the region-based refocusing and deconvolution of fluorescence images provide robust image quality, particularly in those imaging systems with a large FOV where optical aberrations and defocusing effects are more pronounced. In addition, phase image–based stitching and registration leverages label-free quantitative phase images to align images. Since phase images remain stable across imaging cycles, this approach ensures consistency without the need for a dedicated fluorescence reference channel. This not only provides a quantitative mapping of tissue architecture but also increases the number of fluorescence targets that can be imaged per cycle, streamlining the overall workflow.

This pilot study provides a proof-of-concept for the potential utility of this multi-modal image processing platform via quantitative evaluation of mucosal healing in patients with ulcerative colitis. By quantifying the percentages of major cell types, immune cell activation (HLA-DR), cell proliferation (Ki67), and epigenomic states reflecting chromatin compaction and transcriptional activities, and spatial adjacency between different cell types, our platform shows that quantitative metrics can clearly differentiate normal colon tissue, inflamed tissue, and normal-appearing tissue in patients in remission.

Notably, a higher percentage of active immune cells (HLA-DR) and proliferating immune and epithelial cells (Ki67) is a key characteristic of inflamed tissue. These metrics also indicate an intermediate mucosal healing status in patients in remission. Our analysis reveals a higher percentage of cytotoxic T cells and active neutrophils in patients in remission, offering potential insights into the dominant functional immune cells in regulating mucosal healing in remission. Additionally, our image-processing pipeline evaluates global epigenomic states of chromatin compaction and repressive transcription, distinguishing control, inflamed, and remission groups.

Beyond the overall expression levels of functional proteins, our quantitative analysis of spatial adjacency between epithelial cells and immune cells shows that the highest percentage of neutrophils are located in close proximity to epithelial cells in inflamed tissue as well as in some patients with remission. This metrics of neutrophil infiltration into epithelium and lamina propria aligns with histological criteria for assessing disease activity in UC [20]. Further evaluation of the spatial relationship among different cell types could provide some valuable insights into the underlying characteristics of mucosal healing and improved metrics for assessing disease activities in remission.

In addition to serving as a reference for image registration, this study also supports the potential of quantitative phase imaging in assessing epigenomic changes in the cell nuclei linked to inflammation. As illustrated in **Figure 9**, the average phase values of individual nuclei correlate with the degree of chromatin condensation (marked by H3K27me3 and H3K9me3) in both UC patients and controls. Notably, cells with more open, transcriptionally active chromatin states exhibit lower refractive indices (phase values), a trend consistent across immune and epithelial cell populations. These findings suggest that label-free QPI, within this multi-modal imaging platform, could be harnessed in the future to evaluate global chromatin architecture at the single-cell level.

We acknowledge the small sample size of this study, which primarily aims to demonstrate the feasibility of our multi-modal image processing pipeline in quantifying the tissue microenvironment and assessing the epigenomic status of individual cells in mucosal healing. Despite the limited number of patients, we analyzed between 10,000 and 40,000 cells per patient, generating a substantial single-cell dataset that captures considerable cellular heterogeneity. Moving forward, we plan to expand the sample size and conduct a more comprehensive analysis, examining the intestinal epithelium, immune microenvironment, extracellular matrix, and their spatial interactions across a broader patient cohort to further validate the clinical relevance of this multi-modal imaging platform.

## 4 Methods

### 4.1 Data acquisition and online autofocusing

In the image processing pipeline, the quantitative phase image provides quantitative mapping of unbiased tissue architecture via refractive index of different sub-cellular components, which can also be used for multi-channel and multi-cycle image registration and stitching. In our image acquisition workflow, bright-field images and fluorescence images are acquired consecutively. **Supplementary Fig. 5(a)** presents the schematics of the imaging system, where fluorescence images are captured in the reflection mode, and bright-field images in the transmission mode.

When acquiring data, an online autofocusing method was employed to determine the focal plane in the bright-field images. This method is illustrated in **Supplementary Fig. 5(b)**, where autofocusing is achieved through calculating the normalized variance (*NVAR*) [21] of the brightfield image. The *NVAR* is defined by [22]:

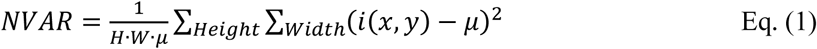

Here in Eq. (1), *H* and *W* denote the height and width of the image, while *μ* represents its mean intensity. This calculation compensates for variations in average intensity across different images by normalizing with *μ*. The optimal focal plane is identified at the point where *NVAR* is minimized [23].

Furthermore, as depicted in **Supplementary Fig. 5(c)**, field of curvature can be observed in the captured images due to the curved surface of the optics [24, 25]. This results in a discrepancy in the focal plane between the central and peripheral areas of the image, particularly pronounced in systems with an extended field of view. For instance, for the microscope system used in our research has a 2mm x 3mm FOV [26]. It exhibits a focal plane discrepancy of approximately 15 *μm* from the center to the edge.

After determining the focal plane, to overcome the field of curvature induced aberration, we capture a set of z-stack sequences at different axial locations in both fluorescence and bright-field images, as demonstrated in **Supplementary Fig. 5(c)**. This information is used subsequently to correct field-of-curvature aberration to ensure sharp focus across the entire FOV, thereby enhancing the accuracy and quality of our imaging data.

### 4.2 Quantitative phase retrieval using region-based transport-of-intensity (TIE) equation

The quantitative phase image at each section is reconstructed by solving TIE [27]:

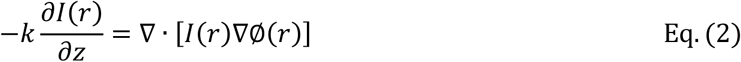

Where *k* and *r* are the wave number and the 2D spatial coordinates, respectively, in Eq. (2). This equation relates the changes in light intensity *I*(*r*) to the phase distribution Ø(*r*) to be solved. The axial intensity derivative on the left-hand side of the equation can be estimated using the two-plane finite-difference method, which is given by [28]:

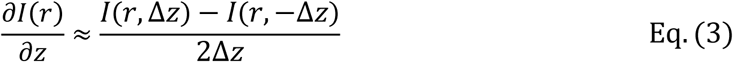

where *I*(*r*, Δ*z*) and *I*(*r*, −Δ*z*) are two slightly defocused images taken at equal but opposite distances from the focal plane in Eq. (3). The quantitative phase image is obtained by solving the equation with a FFT based method [29–31]. In our case, a total of 10 axial positions were acquired.

### 4.3 Flat-field correction of fluorescence image

We compensate for the nonuniform detection efficiency by implementing a calibration map on each detection channel. The process involves several steps to ensure accurate and uniform intensity across the entire fluorescence image.

1. We prepare a sample with a dense distribution of fluorescence markers, such as cell nuclei. Twenty fluorescence images are acquired at different locations to suppress intensity variations of the fluorescent targets.
2. The images are binned with a scaling factor of 0.1 to eliminate structure-induced intensity bias, thus reducing noise and ensuring that the detection efficiency map represents broader trends rather than specific cellular structures.
3. Through spot finding algorithms, the position and intensity of each fluorescence marker in the binned images are determined. These measurements are used to assess the detection efficiency at various positions within the field of view.
4. The calibration map is created to account for the system-induced intensity variation across the entire field of view. To implement such a map, the above steps 2-3 are repeated across the 20 images, and the data are combined to create a calibration map. A cubic polynomial surface fitting is then applied to smooth the detection efficiency pattern, ensuring that the map accurately represents the detection efficiency variations across the field of view.
5. Using the flat-field calibration map, a point-by-point calibration is applied to the fluorescence images. This calibration adjusts the intensity values based on the detected efficiency, compensating for any nonuniform intensity variation induced by the imaging system.
6. After applying the flat-field correction, we achieve a relatively uniform intensity across the entire fluorescence image. This method effectively mitigates the influence of nonuniform detection efficiency, resulting in more accurate and consistent fluorescence imaging.

By following these steps, we ensure that the fluorescence images are corrected for any system-induced non-uniformity, leading to more reliable quantitative results in the imaging experiments.

### 4.4 Background removal of fluorescence image

As illustrated in **Supplementary Figs. 6(a-b),** the Otsu’s threshold method is first used to extract the background from the raw fluorescence image. However, this global thresholding can mistakenly remove targets with weak signals. To enhance the accuracy of the background identification, the histogram of the background (shown in **Supplementary Figs. 6(c)**) is calculated, and the threshold is set at the peak of this histogram. Subsequently, subtracting this threshold results in an image with significantly improved contrast compared to the original raw image, as demonstrated in **Supplementary Fig. 6(d)**.

### 4.5 Region-based deconvolution of the fluorescence image

As illustrated in **Supplementary Figs. 4(a)-(b)**, the original fluorescence image is divided into smaller regions, each corresponding to an area where the PSF was experimentally determined. The size of each region was 300 μm × 300 μm. This is chosen to ensure a uniform PSF to be experimentally measured within each region for accurate deconvolution.

The region-based deconvolution is performed using a two-step method. Firstly, the focal plane of the fluorescence image of a single region is identified by locating the z-stack image with the highest frequency content. Each image in the z-stack is transformed into the frequency domain using a 2D fast Fourier transform (FFT), and the focal plane is determined by comparing the ratio of high to low-frequency components. Then, a high-speed three-dimensional linear deconvolution [6, 7] is applied to the fluorescence image using region-specific PSF.

### 4.6 Determination of the focal plane based on 2D Fourier transform

The sharpest fluorescence image at the focal plane presents a higher ratio between high-frequency and low-frequency components compared to out-of-focus images. As illustrated in **Supplementary Figs. 7(a-b)**, each fluorescence image in the z-stack undergoes 2D FFT to produce its respective 2D spectrum. Subsequently, as shown in **Supplementary Figs. 7(c)**, a round-shaped mask is used to separate the high and low-frequency components, with the mask size determined empirically. Finally, the ratios between the high and low-frequency components are computed, and the focal plane is identified as the image with the highest ratio, as depicted in **Supplementary Figs. 7(d)**.

### 4.7 Multi-tile stitching using quantitative phase images

To seamlessly stitch the two adjacent image tiles, a critical step is to correct the spatial shifts between adjacent tiles [32–34] through image registration of their overlapping regions (approximately 13% here). The shifts between adjacent image tiles are corrected using the phase correlation method [9, 10]. This method computes the phase shift in the frequency domain, which effectively indicates the spatial shift between two images. The peak of the phase correlation with the highest normalized cross-correlation is selected as the translation between two adjacent images.

After correcting the spatial shifts, the whole-slide image is obtained by fusing the image tiles using linear blending method. This method calculates a weighting factor for each pixel in the overlapping area according to the distance to its own border [10, 11], and can effectively reduce the brightness difference in the overlapping area.

Following the stitching of the phase image, the whole-slide fluorescence images for all color channels are generated by applying the same transformation matrix derived from the phase images to the fluorescence images.

### 4.8 Multi-cycle image registration based on quantitative phase images

A set of 4-5 multiplexed fluorescence images are obtained at each imaging cycle and several imaging cycles are often required to obtain a total of 16 markers. To register all images acquired at different imaging cycles, we used phase images acquired at each cycle as the reference and registered all images using the Scale-Invariant Feature Transform (SIFT) method [13–15]. Phase images provide the image contrast of tissue architecture independent of the labeled fluorescence targets. This method extracts distinctive features from images that are invariant to scaling, rotation and illumination, and the feature vectors from different images are matched for registration. To balance the trade-off between accuracy and computational efficiency, the feature threshold was adaptively adjusted to control the number of key feature points within a desirable range for efficient processing.

### 4.9 Multi-model image registration

As shown in **Supplementary Fig. 8(a)**, besides the image registration between phase and fluorescence images, whole-slide histopathology image is often needed as a gold standard for pathological assessment. As the whole-slide histology image is often acquired using standard commercial whole-slide scanner distinct from the imaging systems used for acquiring phase and fluorescence images. Registering multi-modal images is particularly challenging due to two key factors. First, different imaging modalities produce images with varying intensity, contrast, resolution, and pixel size, making it difficult to identify corresponding structures. Second, during the experimental process, the sample may have shrinkage, expansion, or distortion, further complicating alignment. To address these challenges, a series of preprocessing steps are applied to ensure that the images are compatible for further analysis. Firstly, the H&E image, initially a negative image with a bright background and dark tissue signals, is converted to a positive image by subtracting each pixel value from the image’s maximum intensity, as shown in **Supplementary Fig. 8(b)**. Following this, intensity normalization is then performed on all multi-modal images, via dividing their pixel values by the respective maximum intensity to minimize modality-based differences. Next, the orientation of the fluorescence image is adjusted to align with the H&E image, facilitating tissue structure matching. This step improves the normalized cross-correlation coefficient (NCC) from 0.02 to 0.13, as shown in **Supplementary Fig. 8(c)**. Finally, the fluorescence image is resized to match the resolution and pixel size of the H&E image, further increasing the NCC to 0.19.

After these preprocessing steps, the registration of histopathology images with fluorescence images is performed using a mutual information-based registration method [16], as shown in **Supplementary Fig. 8(d)**. This method maximizes the statistical dependence between the images, ensuring precise alignment despite the differences in these modalities, the NCC is increased to 0.67. As a result, multi-modal images, including quantitative phase, fluorescence, and H&E images, are properly registered, and can be used for subsequent correlative analysis.

### 4.10 Voronoi Cell Diagrams and Cell Adjacency Matrices

To create the Voronoi cell diagrams and respective adjacency matrices Cellpose [29, 30] was used to segment cells and create binary masks for each of the five cell type markers of interest (CD4, CD8, CD68, MPO and Cytokeratin) from the raw fluorescence images. For each segmented cell the centroid and bounding box was computed and cells with an area smaller than 4 pixels were treated as artifacts and ignored. For each segmented cell, the cell type was assigned by finding the cell marker binary mask with the largest number of pixels within the respective bounding box. If no marker was present within the bounding box, the cell remained unassigned and was ignored. A Voronoi diagram and respective cell adjacency matrix was computed using the centroids of the assigned cells. Cell adjacencies from the Voronoi diagram with a Euclidian distance between centroids larger than 100 pixels (32µm) were ignored.

### 4.11 Epithelial Cell Adjacency Statistics

The total number of assigned immune cells (Neutrophil, T Killer, Macrophage, T Helper) for each respective specimen with first degree adjacencies to epithelial cells were computed using the cell adjacency matrix as described in Section 4.8. The total counts were per immune cell type were subsequently normalized over the number of epithelial cells.

### 4.12 Patient sample collection

This study was approved by the Institutional Review Board of the University of Pittsburgh. We prospectively recruited 6 patients with ulcerative colitis and 3 controls with no known gastrointestinal disorders undergoing routine screening colonoscopies at the University of Pittsburgh Medical Center. The rectal biopsies were obtained and immediately fixed in 10% formalin and then subsequently processed with a standard tissue paraffin embedding. For multiplexed imaging, we sectioned the tissue at 3 µm, placed onto a #1.5 coverslip, which is mounted on an imaging chamber for multi-cycle phase and fluorescence imaging.

### 4.13 Preparation of tissue section

The preparation and imaging of the UC sample follow a systematic procedure outlined as follows: Initially, quantitative phase image is captured images following the standard paraffin removal and rehydration using graded alcohol prior to antigen retrieval. Subsequently, antigen retrieval is carried out by placing the tissue section on a glass slide within a chamber, exposing it to LED light at room temperature for 15 minutes while adding 1mL of bleaching solution (H_2_O_2_, 30% (wt/vol), 0.02M NaOH) into the chamber, followed by rinsing with PBS to remove the bleaching solution. This step is to bleach the tissue autofluorescence.

Following this, permeabilization and blocking are executed by incubating the sample with 0.2% Triton X-100 in PBS, followed by blocking with 4i blocking solution (sBS, 1% BSA, 100mM NH_4_Cl, 150mM Maleimide) for 30-60 minutes [35]. Immunofluorescence staining is subsequently carried out by incubating the primary antibody either for 2 hours at room temperature or overnight at 4°C, followed by washing with PBS and incubating with the secondary antibody for 1 hour, with additional PBS washes. During fluorescence imaging, imaging buffer (700mM NAC in ddH_2_O, adjusted to pH 7.4) is added to the tissue sample to prevent photo-crosslinking. Antibody elution follows each cycle of fluorescence imaging, involving washing with PBS and incubating the sample with elution buffer (EB, 0.625M L-Glycine, 3.75M Urea, 3.75M GC in ddH_2_O, adjusted to pH 2.5) and TCEP solution (350mM TCEP in ddH_2_O) for 15 minutes, repeated three times. The chemicals used are listed in **Table 1** and the antibodies used for the multi-cycle fluorescence imaging are listed in **Table 2**. This multi-cycle fluorescence imaging process is repeated until all imaging cycles are completed.

**Table 1.** Chemicals used for tissue section preparation.

|  | **Name** | **Manufacturer** | **Item number** |
| --- | --- | --- | --- |
| 1 | Xylene | Fisher Scientific | X3P-1GAL |
| 2 | Ethanol (100%) | Fisher Scientific | BP28184 |
| 3 | 10X Antigen retrieval buffer (PH6) | Invitrogen | 00-4955-58 |
| 4 | Sodium borohydride | Sigma-Aldrich | 213462-25G |
| 5 | Triton X-100 |  |  |
| 6 | Hydrogen peroxide 30% W/W | Sigma-Aldrich | 216763-100ml |
| 7 | Sodium hydroxide, Pellet AR (ACS) | Mallinckrodt Chemicals | 7708-10 |
| 8 | 10X PBS with calcium and magnesium | Fisher Scientific | BP399 |
| 9 | L-Glycine | Sigma-Aldrich | 50046-50G |
| 10 | Urea | Sigma-Aldrich | U5378-500G |
| 11 | Guanidium chloride (GC) | Millipore-Sigma | 369075-1KG |
| 12 | N-acetyl-cysteine (NAC) | Sigma-Aldrich | A7250-100G |
| 13 | Ammonium chloride(NH4Cl) |  |  |
| 14 | Tris(2-carboxyethyl)phosphine hydrochloride (TCEP) | Sigma-Aldrich | C4706-10G |
| 15 | Maleimide | Sigma-Aldrich | 129585-2G |
| 16 | 12M Hydrochloric acid | Fisher Scientific | A144-500 |
| 17 | Trizma base | Sigma-Aldrich | T1503-25KG |
| 18 | Bovine serum albumin | Sigma-Aldrich | A9647-100G |

**Table 2.** Antibody used for multi-cycle fluorescence imaging.

| Antibody | Catalog number | Dilution | Conjugated Dye | Secondary antibody | Cycle |
| --- | --- | --- | --- | --- | --- |
| CD4 | ab288724 | 1:100 | Alexa488 |  | 1 |
| CD19 | 396304 | 1:200 | Alexa647 |  | 1 |
| HLA-Dr | Ab92511 | 1:200 | Unconjugated | Goat anti-rabbit-CF532 | 1 |
| TNFα | NBP2-45331 | 1:50 | Unconjugated | Goat anti-mouse-Alexa594 | 1 |
| H3K9me3 | Ab8898 | 1:200 | Alexa647 |  | 2 |
| CD68 | Ab213363 | 1:200 | Unconjugated | Goat anti-rabbit-CF532 | 2 |
| CD16 | 302007 | 1:100 | PE |  | 2 |
| Ki67 | 9449S | 1:200 | Unconjugated | Goat anti-mouse-Alexa594 | 2 |
| H3K4me3 | 12064S | 1:200 | Alexa647 |  | 3 |
| CD8 | Ab237709 | 1:400 | Unconjugated | Goat anti-rabbit-Alexa594 | 3 |
| αSMA | SC-20060 | 1:1000 | Alexa488 | | 3 |
| FoxP3 | 58-5773-82 | 1:100 | Alexa532 |  | 3 |
| H3K27me3 | 12158S | 1:500 | Alexa647 |  | 4 |
| Cytokeratin | 53-9003-82 | 1:3000 | Alexa488 |  | 4 |
| MPO | af3667 | 1:200 | Unconjugated | Donkey anti-goat, Alexa594 | 4 |
| CD163 | SC-33715 | 1:100 | Alexa546 |  | 4 |

### 4.14 Quantitative analysis of the UC tissue

To analyze the UC tissue, we initiated our process with the segmentation of cells using Cellpose [36, 37], as shown in **Supplementary Figs. 9(a)**. This segmentation process generates cell masks containing the location and area information of each cell within the sample. Red blood cells (RBCs) present a unique challenge in this context. Typically, RBCs exhibit relatively high signal intensity within the 400 nm to 600 nm wavelength channels, making them difficult to identify in fluorescence images. However, they are readily identifiable in H&E-stained histology images due to their distinct staining characteristics. Leveraging this, we create a specific mask for RBCs using the information from the H&E images, as depicted in **Supplementary Figs. 9(b)**. This RBC mask is then used to refine our initial cell mask by removing the identified RBCs, resulting in a final mask that exclusively represents the cells of interest.

To determine the thresholds used for defining cell types, a 3-point moving average filter was first applied to the fluorescence image to smooth out noise and enhance the signal. Following this, the cytokeratin image was subtracted from the fluorescence image to eliminate any potential crosstalk interference. Afterwards, a representative background area was manually selected. The mean intensity value of this selected background was then calculated and utilized as the threshold to extract the signal regions in the image.

After removing the background, to quantitatively analyze the fluorescence images, we start by binarizing each image and then overlaying it with a cell mask for comparison, as demonstrated in **Supplementary Figs. 9(c)**. Within each cell’s defined area as per the mask, we quantify the cell properties such as the number of cells, and fluorescence signal intensities (**Supplementary Figs. 9(d)**). These measurements allow us to assess the expression and localization of specific markers within individual cells.

## 6 Author contributions

**MC** developed the image processing pipeline, analyzed the experimental data, and drafted the original manuscript. **HM** designed the multi-modal microscope imaging system, developed the flat-field correction, calculated the system resolution, and provided guidance on the development of the image processing pipeline. **XS, JX, and PN** designed biological experiments and conducted imaging experiments. **DSG** contributed to data analysis. **MS**, **REB**, **LS**, **BAM**, **RMB** coordinated clinical studies, recruited the patients, collected the patient biopsies and identified the clinical information. **YL** provided overall guidance on the study design, helped write the original draft and revise the manuscript, and provided funding support for this work. All authors read and approved the final manuscript.

## 7 Acknowledgements

We acknowledge the assistance of Chaojie Zhang in calculating the Fourier Ring Correlation based resolution of the images. This study was funded by the National Institute of Health grants R01CA254112, R01CA232593, and P41EB031772.

## 9 Data availability

The example datasets that accompany the code are available in the Zenodo repository: (https://doi.org/10.5281/zenodo.14838554).

## 10 Code availability

The code for the data processing pipeline presented in this study is available in Github and can be accessed via this link: (https://github.com/YangLiuLab/MM_WSI).

