## Supplementary Figures for "A multi-modal whole-slide image processing pipeline for quantitative mapping of tissue architecture, histopathology, and tissue microenvironment"

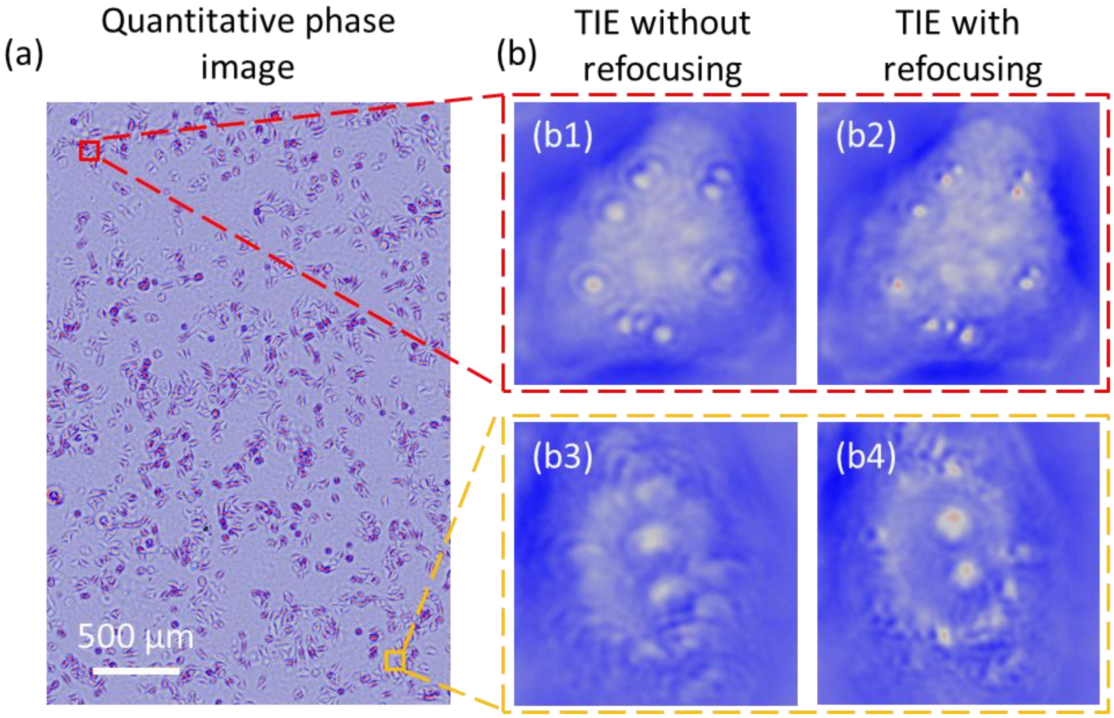


**Supplementary Fig. 1 The performance of reconstructed quantitative phase image. (a)** The quantitative phase image of the entire FOV. **(b)** The phase images of individual cells on the corner areas. Region-based TIE (**b1-b2**) shows a better resolution compared to the traditional method (**b3-b4**).

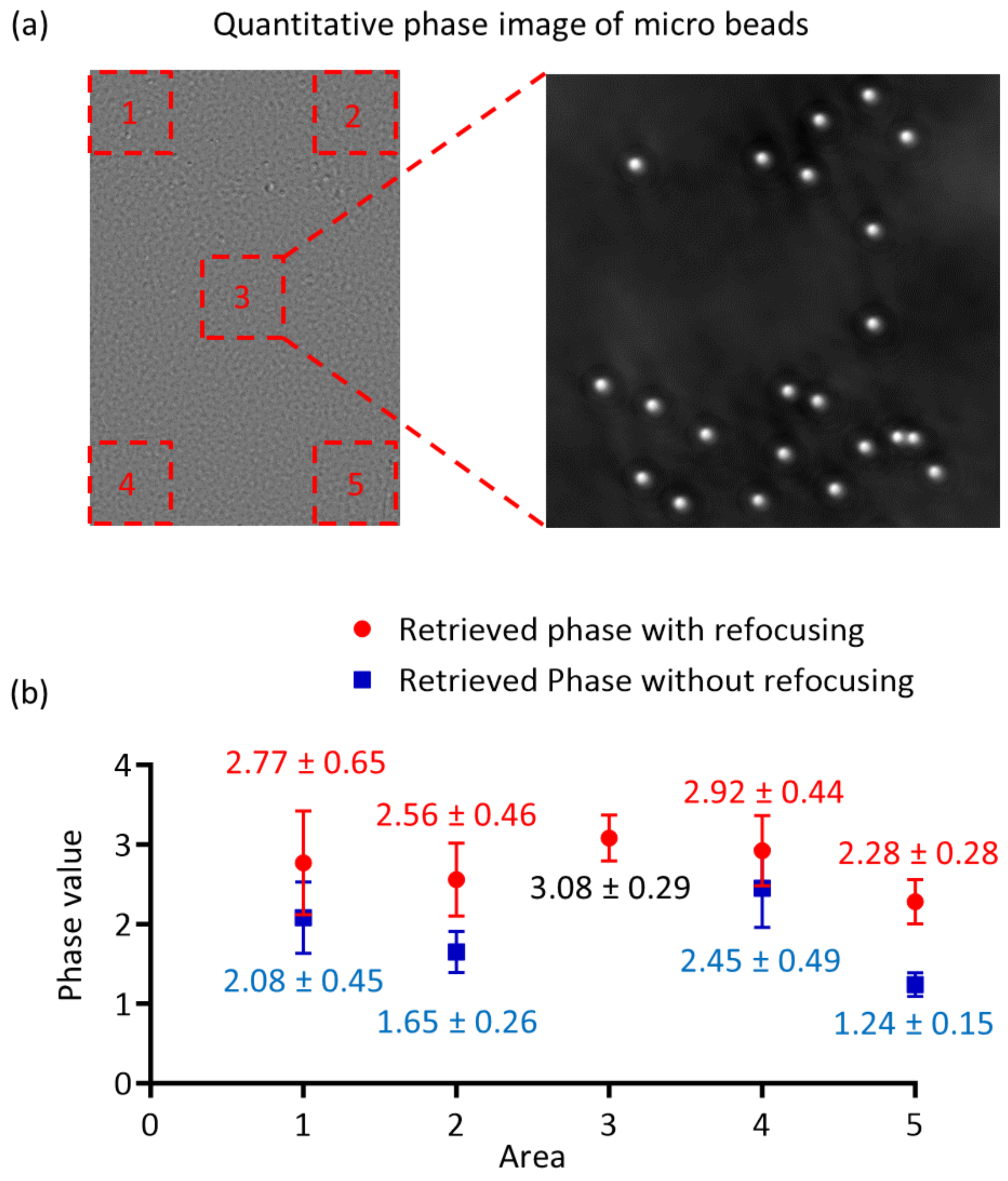


**Supplementary Fig. 2 Phase values measured using 2μm microspheres. (a)** The quantitative phase image of polystyrene microspheres. Five areas covering the four corners and the center area are selected to estimate the accuracy of the phase values of the microspheres. **(b)** The comparison of the phase values was calculated using TIE-based phase retrieval with and without refocusing. The 2 μm polystyrene microspheres are immersed in glycerol. The refractive indexes of the microspheres and the medium are 1.59 and 1.47, respectively. Thus, the expected phase value of the microsphere is 2.96. Using TIE-based phase retrieval with refocusing, the retrieved phase values at the corner areas are more accurate compared to the results without refocusing.

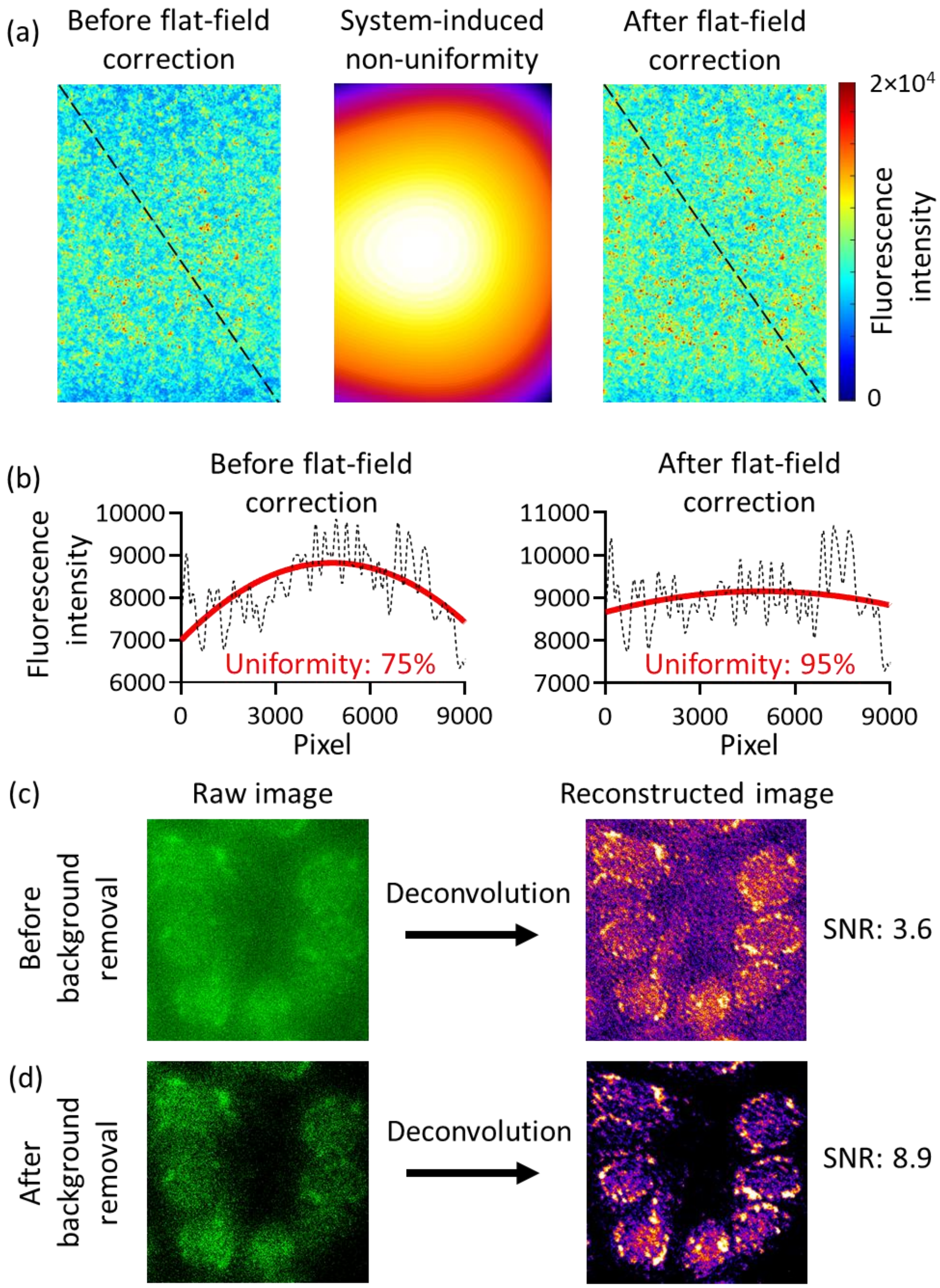


**Supplementary Fig. 3 Preprocessing of fluorescence images.** (**a**) Flat-field correction is performed by compensating the fluorescence image with a calibration map. The flat-field corrected image shows better intensity uniformity across the entire image. (**b**) The quantified uniformity of the fluorescence images. The black curve is the intensity profile along the black dashed lines in the fluorescence images in (a), and the red solid lines are fitted from the profiles. After flat-field correction, the uniformity improves from 75% to 95%. (**c-d**) The raw and reconstructed fluorescence images of the cells stained for Caspase-3 (c) before and (d) after background noise removal. The SNR was increased from 3.6 to 8.9.

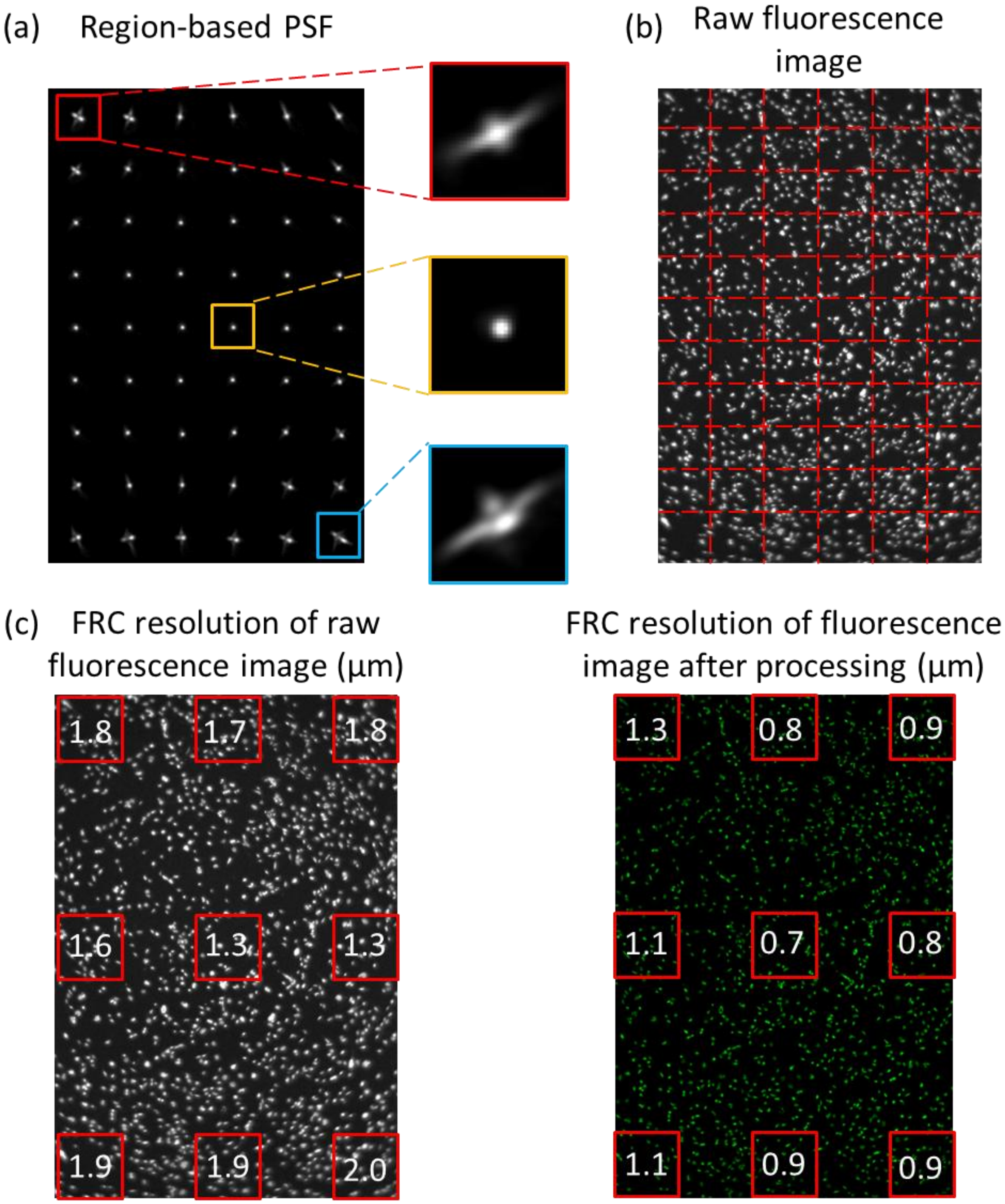


**Supplementary Fig. 4 Region-based PSFs and deconvolution. (a)** PSFs vary across the entire FOV, based on the measurement of fluorescence microspheres. The PSFs in the peripheral areas exhibit serious distortions. **(b)** The raw fluorescence image of SW480 cells fluorescently stained for centromeres, which is divided into small regions corresponding to the PSFs. **(c)** The Fourier Ring Correlation (FRC) resolution of the fluorescence images before and after processing. It shows an average of approximately 40% improvement in FRC resolution after processing.

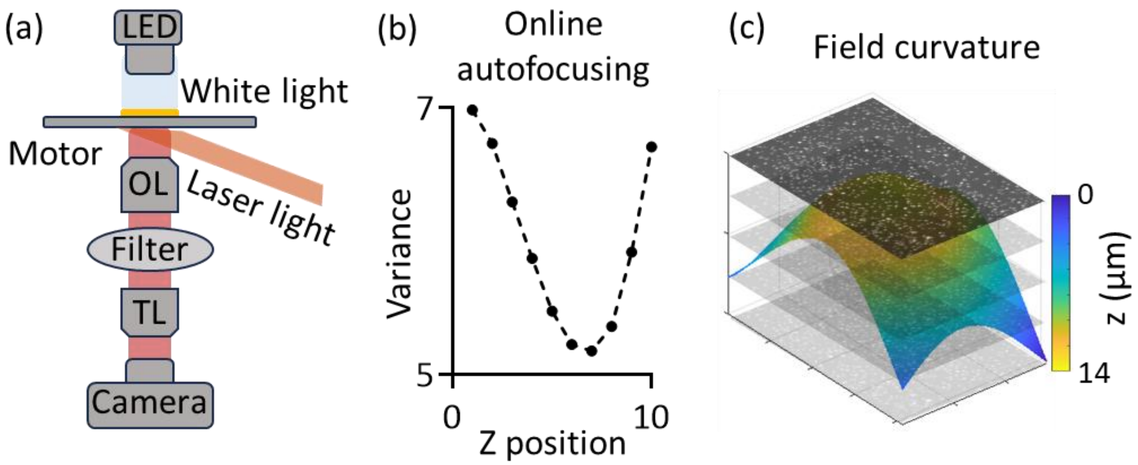


**Supplementary Fig. 5 Data acquisition and online autofocusing. (a)** Schematics of the imaging system. The fluorescence imaging is in the reflection mode and the phase imaging is working in the transmission mode. OL: objective lens, TL: tube lens. **(b)** Online autofocusing scheme by minimizing the normalized variance of the bright-field image. **(c)** The field of curvature.

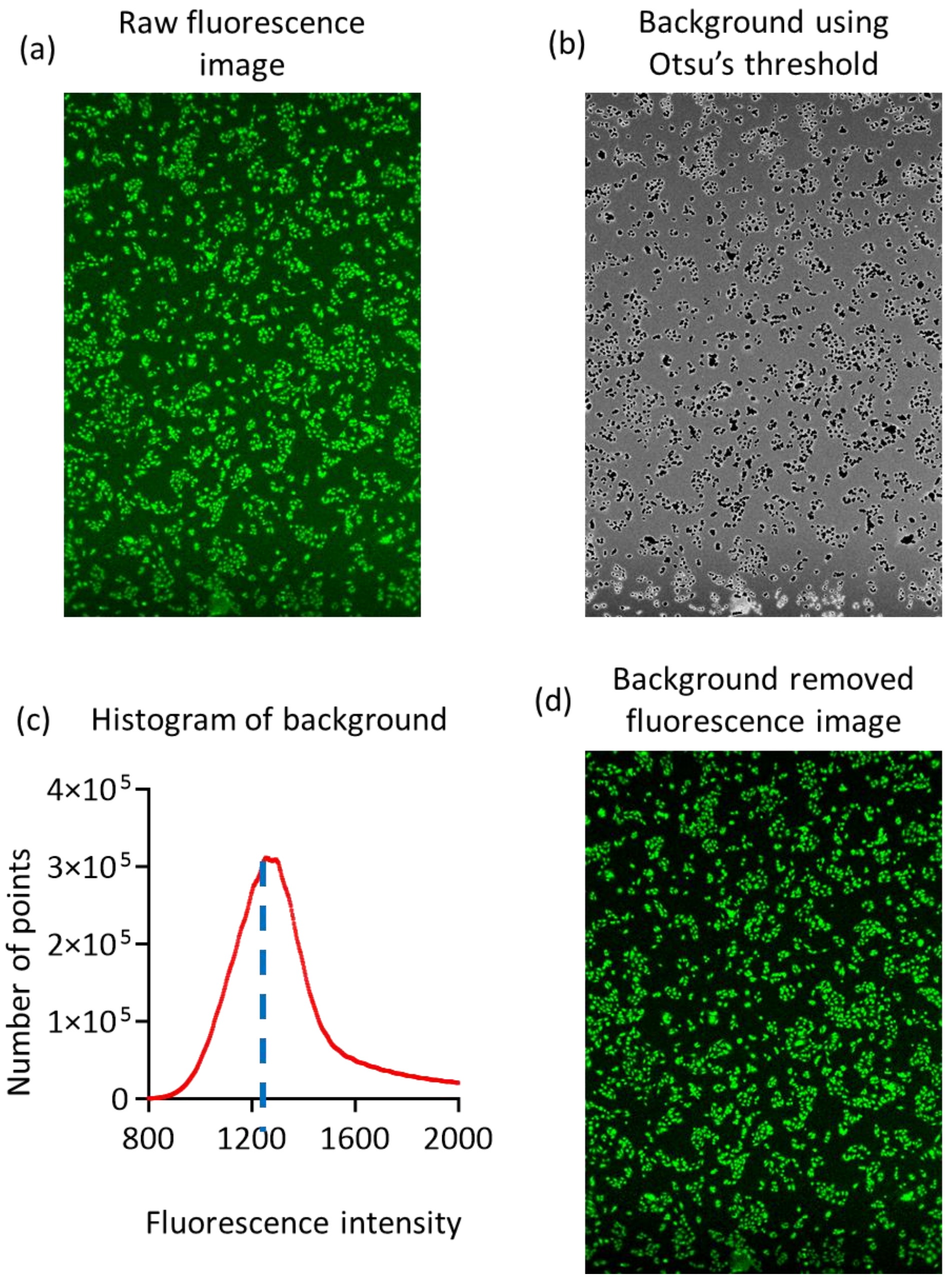


**Supplementary Fig. 6 Background removal of the fluorescence image. (a)** Raw fluorescence image. (**b**) The background extracted from the raw fluorescence image using the threshold calculated by Otsu's method. (**c**) The histogram distribution of the background signal. The final threshold is determined by the peak value of the histogram distribution of background fluorescence intensity. **(d)** The fluorescence image after background removal.

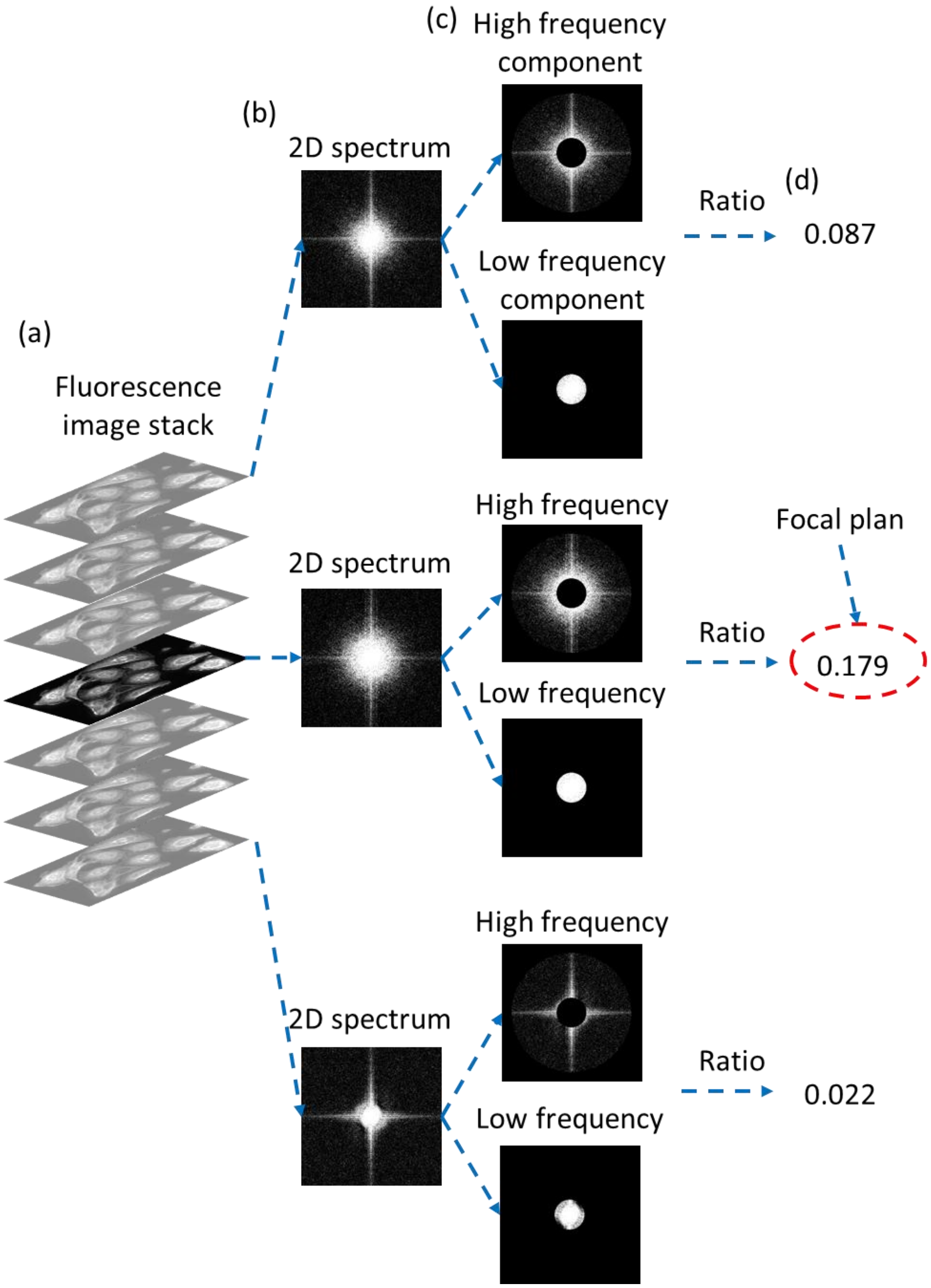


**Supplementary Fig. 7 Determine the focal plane of the fluorescence image. (a)** The raw fluorescence image stack. **(b)** Calculate the 2D Fourier spectrum of all the images. **(c)** Division of the 2D spectrum into high-frequency and low-frequency components. **(d)** Calculation of the ratio between the high and low-frequency components of the 2D Fourier spectrum. The focal plane is defined as the image with the highest ratio.

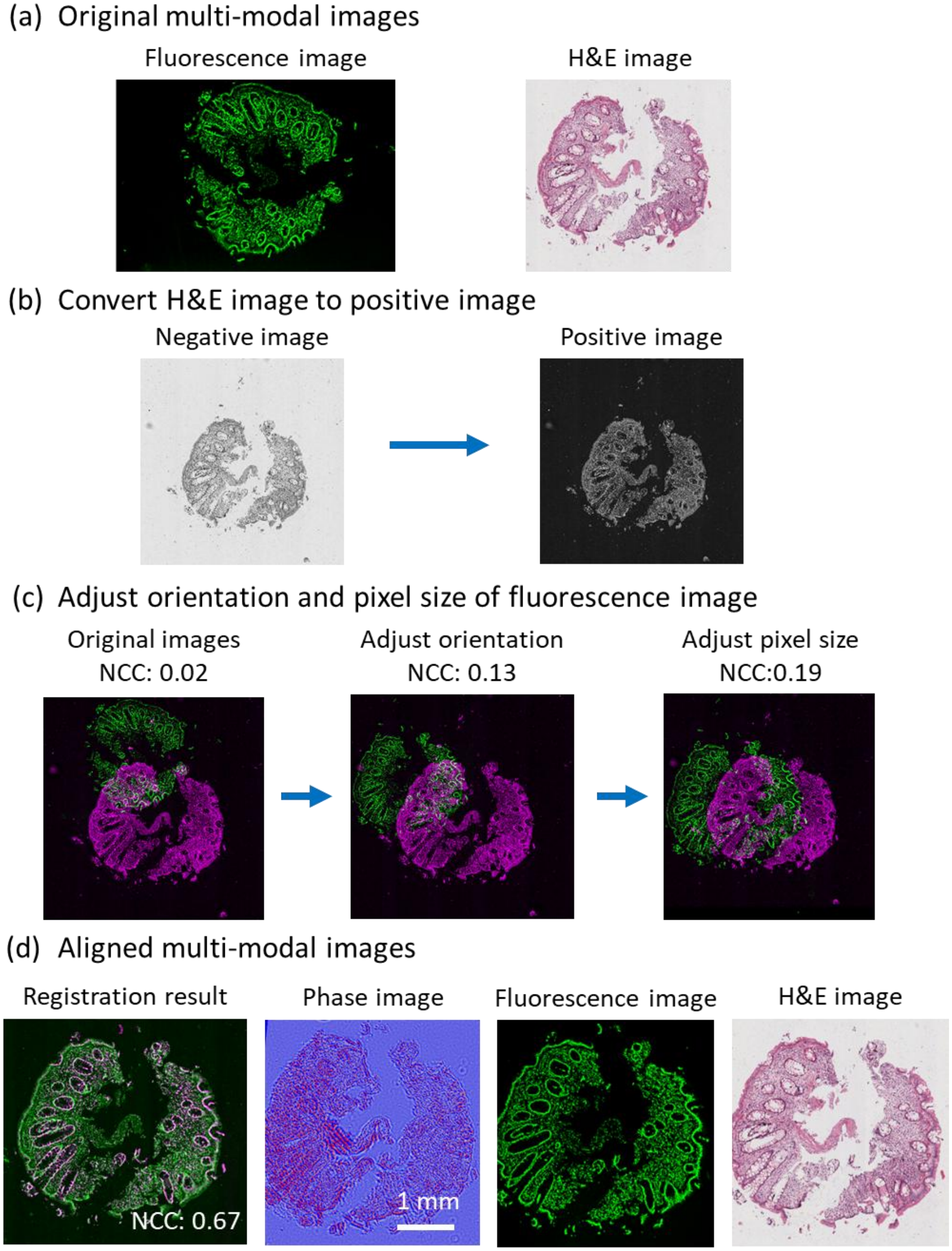


**Supplementary Fig. 8 Procedure of multi-modal registration. (a)** The original multi-modal images. The fluorescence image is stained with H3K9me3. **(b)** Conversion of the H&E image from a negative to a positive image. **(c)** Adjusting the orientation and pixel size of the fluorescence image to match with the H&E image. The normalized cross-correlation coefficient (NCC) between the two images is increased from 0.02 to 0.19 after the steps. **(d)** The registration results and the aligned multi-modal images. The NCC is further improved to 0.62 after the mutual information-based registration.

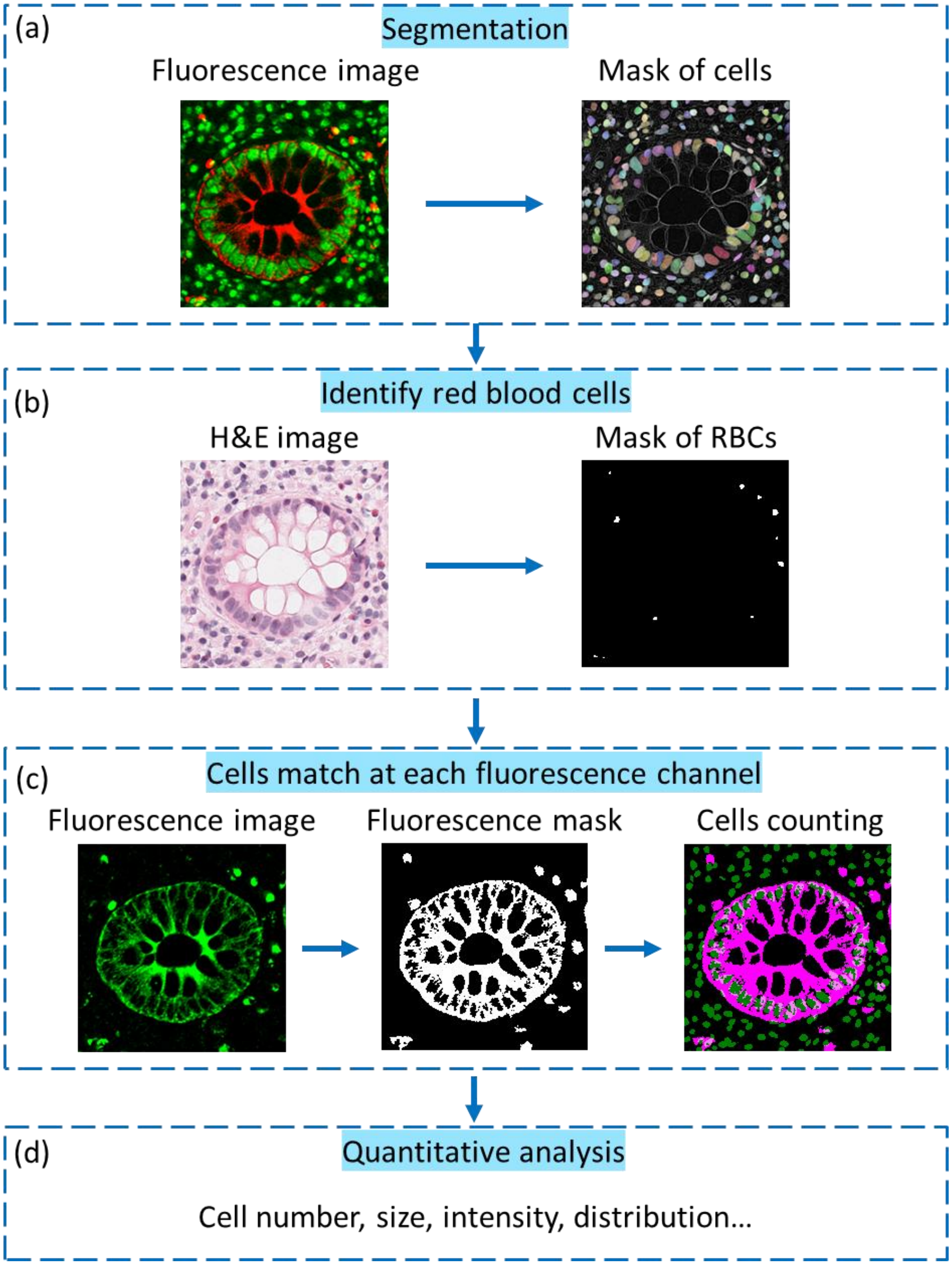


**Supplementary Fig. 9 Quantitative analysis of UC tissue. (a)** The cells are segmented using Cellpose and cell masks are generated from the fluorescence image. **(b)** The mask of red blood cells generated from the H&E-stained histology image. It is then subtracted from the cells mask. **(c)** The cells in each fluorescence channel are determined by comparing the binarized fluorescence image with the cells mask. **(d)** Quantitative analysis of the segmented cells.
